# Aberrant expression of histone H2B variants reshape chromatin and alter oncogenic gene expression programs

**DOI:** 10.1101/2024.11.18.624207

**Authors:** Wesley N. Saintilnord, Youssef A. Hegazy, Kristin Chesnutt, Meredith Eckstein, Richard N. Cassidy, Héjer Dhahri, Richard L. Bennett, Daniёl P. Melters, Elisson Lopes, Zhen Fu, Kin Lau, Darrell P. Chandler, Michael G. Poirier, Yamini Dalal, Jonathan D. Licht, Yvonne Fondufe-Mittendorf

## Abstract

Chromatin architecture governs DNA accessibility and gene expression. Thus, any perturbations to chromatin can significantly alter gene expression programs and promote disease. Prior studies demonstrate that every amino acid in a histone is functionally significant, and that even a single amino acid substitution can drive specific cancers. We previously observed that naturally occurring H2B variants are dysregulated during the epithelial to mesenchymal transition (EMT) in bronchial epithelial cells. Naturally occurring H2B variants differ from canonical H2B by only a few amino acids, yet single amino acid changes in other histone variants (e.g., H3.3) can drive cancer. We therefore hypothesized that H2B variants might function like oncohistones, and investigated how they modify chromatin architecture, dynamics, and function. We find that H2B variants are frequently dysregulated in many cancers, and correlate with patient prognosis. Despite high sequence similarity, mutations in each H2B variant tend to occur at specific “hotspots” in cancer. Some H2B variants cause tighter DNA wrapping around nucleosomes, leading to more compact chromatin structures and reduced transcription factor accessibility to nucleosomal DNA. They also altered genome-wide accessibility to oncogenic regulatory elements and genes, with concomitant changes in oncogenic gene expression programs. Although we did not observe changes in cell proliferation or migration in *vitro*, our Gene Ontology (GO) analyses of ATAC-seq peaks and RNA-seq data indicated significant changes in oncogenic pathways. These findings suggest that H2B variants may influence early-stage, cancer-associated regulatory mechanisms, potentially setting the stage for oncogenesis later on. Thus, H2B variant expression could serve as an early cancer biomarker, and H2B variants might be novel therapeutic targets.

## INTRODUCTION

Chromatin structure dynamically sequesters genomic DNA, providing the framework that regulates gene activity [1, 2]. The nucleosome is the fundamental unit of chromatin and consists of an octamer of four core histones wrapped by ∼147bp of DNA [3–5]. Nucleosome assembly involves sequential deposition of the (H3-H4)_2_ tetramer and H2A-H2B dimers onto DNA, with optional linker histone H1 binding at the nucleosome entry-exit DNA site to further compact chromatin [4].

Disrupting chromatin structure leads to altered pathological gene expression programs, and is a hallmark of cancer [6]. Early studies primarily focused on histone H3 tail mutations because they alter histone post-translational modifications (PTMs) that regulate chromatin structure, access to DNA, and transcription. For instance, H3K27M and G34R/V mutations are linked to pediatric high-grade gliomas [7, 8], while H3K36M and G34RW/L are seen in chondroblastoma and giant cell tumors of the bone, respectively [9–12]. The H3K36M mutation is also implicated in head and neck cancers [13]. Cancer-driving mutations have also been reported in other histones. For example, H1 mutations drive diffuse large B-cell lymphomas [14], while the H2BE76K mutation reduces nucleosome stability and co-occurs with the *PI3KCA* oncogene to enhance cellular transformation [15]. From these and other studies, it has become increasingly evident that many amino acids in a histone are functionally significant, and that even a single amino acid substitution can drive specific cancers, which has led to the concept of an “oncohistone” [10, 16–19].

Chromatin architecture can also be perturbed when cells incorporate naturally occurring histone variants into nucleosomes. Unlike canonical histones, which are expressed specifically in a replication-dependent manner during the S phase of the cell cycle and encoded by multiple genes, histone variants (sometimes referred to as isoforms of the canonical histones) are typically encoded by a single gene and are incorporated into chromatin (and replace canonical histones) throughout the cell cycle [20, 21], independent of DNA replication. These histone variants can differ from canonical histones by only a few amino acids or by significant structural variations. Even the small differences can have important implications for transcription, chromosome segregation, DNA repair and recombination, chromatin remodeling, ADP-ribosylation, germline-specific DNA packaging and activation, and extra-nuclear acrosomal function [20, 22–24]. Most prior studies of histone variants centered on H2A and H3 variants, and revealed that histone variants (like histone mutants) can harbor distinct PTMs that recruit specific chromatin remodelers, thus changing chromatin structure, function, and dynamics [23, 25–27].

Given that histone variants also alter chromatin integrity, it is not surprising that histone variant genes (and their regulators) are also mutated in cancers [28–30], with their occurrence correlating with metastasis and/or survival [16–19]. For example, mutations in the H3.3 variant are associated with pediatric glioblastoma and bone tumors [7, 31], and H2A.X dysregulation promotes the epithelial-to-mesenchymal transition (EMT) in lung cancer [32]. Mechanistically, histone variants sometimes alter oncogene or tumor suppressor gene expression through their specific occupancy at these sites. For instance, reduced H2A.X occupancy at the promoters of EMT transcription factors *SLUG* and *ZEB1* opens chromatin structure and increases their expression [32]. Similarly, the H2A.Z enrichment at EMT gene promoters activates cell cycle regulators and oncogenes in colorectal and breast cancers [33, 34]. On the other hand, macroH2A.1 and macroH2A.2 act as tumor suppressors, silencing EMT transcription factor genes [1, 30, 35].

Environmental toxicogenesis profoundly affects the epigenome, leading to significant health implications including cancer. Previously, we found that H2B protein expression is dysregulated when BEAS-2B cells (derived from normal human bronchial epithelium) undergo inorganic arsenic (iAs)-mediated EMT [35], with specific proteins becoming more abundant. Five genes encode for the same H2B protein whereas others with few amino acid differences are encoded by a single gene [21, 36]. We refer to these as H2B variants to emphasize their single-gene origin, in contrast to canonical H2B, which is encoded by multiple genes. We hypothesized that genome-wide shifts in nucleosome composition resulting from particular H2B variant expression could alter chromatin function and contribute to oncogenesis. Given this intriguing possibility, we investigated how H2B variants modify chromatin architecture, dynamics, and function. We first show that H2B variants are dysregulated in many cancers, and in a cancer type-specific manner. We also found distinct mutational hotspots within several H2B variants, even though H2B variants have DNA and amino acid sequence similarity. We then used biochemical, biophysical, and *in vitro* and cellular assays to determine whether and how H2B variants alter nucleosome physical properties. Altered nucleosome properties translated into altered chromatin accessibility, gene expression, and oncogenic signatures, even without overt *in vitro* oncogenic phenotypes. Based on these findings, we posit that aberrant expression of particular H2B variants may influence early-stage, cancer-associated regulatory mechanisms and “prime” cells for transformation by a future insult. These data indicate that H2B variants shape chromatin architecture, and suggest that they might serve as useful disease-associated biomarkers or targets for therapeutic development.

## RESULTS

### Dysregulation of H2B variants in various cancers is correlated with patient prognosis

Given that genes encoding histone variants are altered in cancer [1, 16, 19, 37], and that H2B variant proteins differ from the canonical and from each other by only a few amino acids (**Fig 1A**), we postulated that H2B variants might also be differentially expressed across cancers. We therefore analyzed RNA-seq data from The Cancer Genome Atlas (TCGA) [38], and found that many H2B variants are indeed dysregulated across various cancers (**Fig 1B**) and in a tumor-specific manner (**Suppl Fig 1A**). To verify that H2B variant expression patterns were not due to broad histone gene regulation, we normalized H2B variant expression to the expression of one of the canonical H2B genes (*H2BC4*) for each cancer. Even after normalization, we still observed significant and cancer-specific differences in H2B variant expression (**Suppl Fig 1B**).

**Fig 1.**
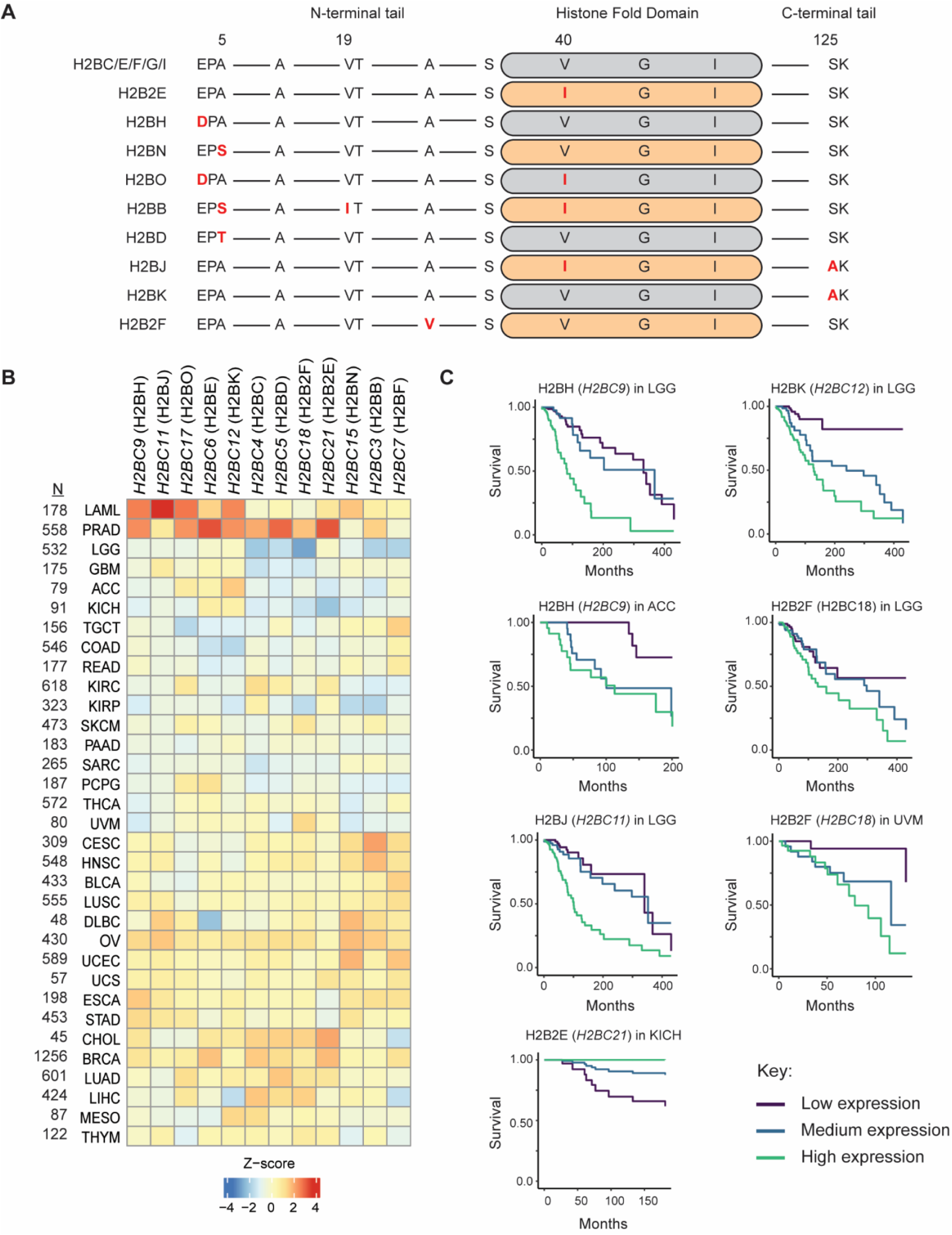
H2B variants are expressed in many cancers, and are associated with patient prognosis. **A**) Amino acid sequence alignment of canonical H2B (top) and H2B variants. Amino acid differences are highlighted in red. **B)** Median expression level of H2B genes across cancer types in The Cancer Genome Atlas (TCGA) database. Columns are H2B variant genes, rows are the cancer type, and N = the number of TCGA patients with the specific cancer type. Protein names are in parentheses. Z-scores are based on variance stabilizing transformed (VST) counts. **C**) Kaplan-Meier 5-year survival curves for TCGA patients and cancers where 5-year survival is significantly associated with H2B variant gene expression. Low expression = below the 33^rd^ percentile; medium expression = between the 33^rd^-67^th^ percentile; high expression = above 67^th^ percentile. Significance of survival risks was determined using the Benjamini-Hochberg false discovery rate (FDR) with a threshold of < 0.05. Only samples with a tumor purity score > 0.7 and library size > 15 million reads were used in the analysis. LGG = low grade glioma; ACC = adenoid cystic carcinoma; UVM = uveal melanoma.

These data prompted us to inquire about H2B variant dysregulation and cancer prognoses. We used the Cox proportional hazards regression model to estimate the association between H2B variant gene expression and 5-year survival for the cancer types and variants where the TCGA database had at least 30 samples. Up-regulation of four H2B variants (*H2BC9, 11, 12* and *18*) but downregulation of one H2B variant (*H2BC21*) that encode H2BH, H2BJ, H2BK, H2B2F, and H2B2E respectively was significantly associated with decreased 5-year survival in four cancers (low grade glioma, adenoid cystic carcinoma, uveal melanoma, and kidney chromophobe; **Fig 1C**), and with age (**Suppl Fig 1C**). These associations were also variant and cancer-type specific. For example, low grade glioma patients with high *H2BC9* (H2BH) expression have a worse prognosis than adenoid cystic carcinoma patients with high *H2BC9* expression. Additionally, while low grade glioma patients who over-express *H2BC11* and *H2BC12* have similar poor survival outcomes, low *H2BC12* expression is associated with a better 5-year survival rate than low *H2BC11* expression (**Fig 1C**).

### Oncogenes and chromatin remodelers are co-expressed with those H2B variants that are associated with poor prognosis

Histone variants could drive cancer directly or exert their influence indirectly through other (non-histone) genes and gene expression programs. To distinguish between these two possibilities, we performed co-expression analysis on the histone genes and cancers described in **Fig 1C**. We found several correlations between H2B variant expression and known oncogenes/chromatin remodelers (e.g., MYC and EZH2; **Suppl Table 1**; **Suppl Fig 2**) and oncogenic pathways (e.g., EGFR, KRAS and STK33; **Suppl Fig 3, 4**). Together, these data suggest that specific H2B variants are associated with specific cancers, and that they contribute to cancer by altering gene expression at multiple loci (to include oncogenes and chromatin remodelers).

### H2B variants exhibit unique mutational hotspots

Specific “hotspot” mutations in histone variants are linked to carcinogenesis [7, 12, 39–41]. We therefore asked if H2B variants also exhibit mutation hotspots. We analyzed a curated set of 205 nonredundant cancer studies consisting of 65,489 sequenced patient samples to identify frequently occurring H2B missense mutations (z score >2). We binned all H2B variants and found the same H2BE76K/Q hotspot we identified previously [15] (**Suppl Fig 5, upper left**). We then parsed the data by H2B variant, and found striking differences in the number and location of hotspot mutations (**Fig 2**; **Suppl Fig 5**). We also found that these hotspot mutations occurred in specific cancers, and differed by H2B variants (**Table 1**). For example, the top hotspot mutation in H2BC (E76K) primarily occurs in bladder cancer, while the top hotspot mutation in H2BJ (G53S/D) is primarily found in non-small cell lung cancer (**Table 1**, **Fig 2**). Although the canonical H2BC protein is encoded by multiple genes (including *H2BC4*, *H2BC6*, and *H2BC7*), we observed distinct hotspot mutations in each *H2BC* gene, each linked to different cancers (**Fig 2**, **Suppl Fig 5**). While gene redundancy likely serves an evolutionary purpose to maintain essential functions, these mutations and the resulting amino acid changes suggest unique impacts on protein function and cancer development. Altogether, several of these mutated variants may weaken histone-DNA interactions, alter chromatin structure, and increase gene transcription in various cancers [9, 10, 42–47].

**Fig 2.**
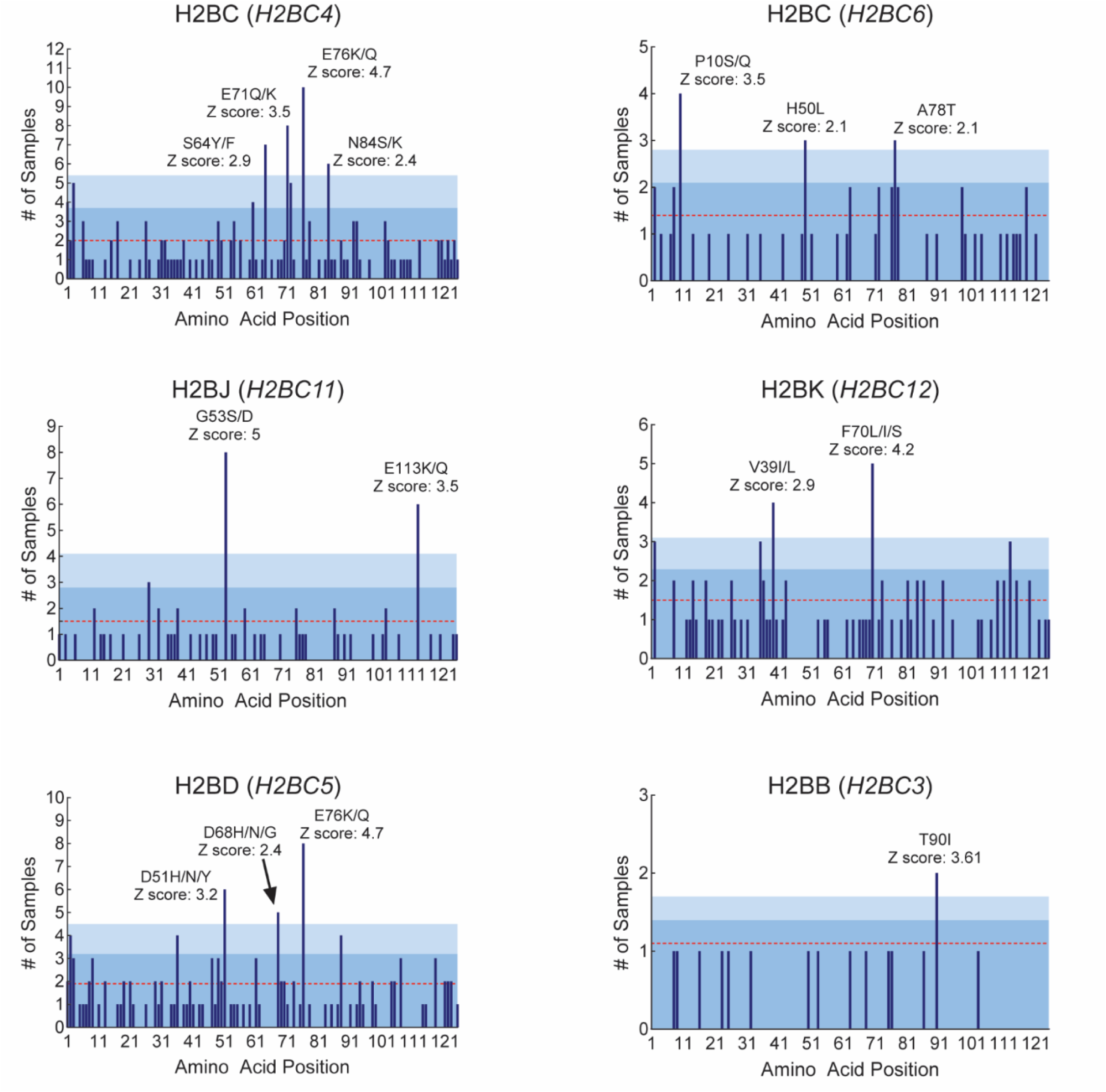
H2B variants exhibit unique mutational hotspots. H2B missense mutations were derived from 65,489 non-redundant patient samples in the cBioPortal for Cancer Genomics database, and encompass all cancers. Frequently occurring mutations are those with a z-score >2, and are labeled in each panel. The gene encoding each H2B variant is indicated in parentheses. Red lines are the average number of mutations for each histone variant, with light blue shading representing the first and second standard deviations from the average, respectively. Notice how canonical H2BC has a different mutational signature depending on which gene is encoding the histone (*H2BC4* and *H2BC6*, respectively). Hotspots for other H2B variants are shown in **Suppl** Fig 5.

**Table 1.** Histone H2B variants hotspot mutations are associated with specific cancers.

| Histones | Hotspot Mutations | Cancers |
| --- | --- | --- |
| H2BC (*H2BC4*) | S64Y/F<br>E71K/Q<br>E76K/Q<br>N84S/K | Colorectal (2), Endometrial (2), Esophagogastric (1), Melanoma (1), Pheochromocytoma (1)<br>Breast (3), Melanoma (2), Non-Small Cell Lung (2), Bladder (1)<br>Bladder (4), Non-Small Cell Lung (2), Cervical (1), Mature B Cell Neoplasm (1), Ovarian (1)<br>Mature B Cell Neoplasm (5), Endometrial (1) |
| H2BJ (*H2BC11*) | G53S/D<br>E113K/Q | Non-Small Cell Lung (5), Bladder (1), Prostate (1)<br>Ampullary (1), Bladder (1), Breast (1), Colorectal (1), Melanoma (1), Squamous Cell (1) |
| H2BB (*H2BC3*) | T90I | Melanoma (2) |
| H2BK (*H2BC12*) | V39I/L<br>F70L/I/S | Bone (1), Breast (1), Endometrial (1), Esophagogastric (1)<br>Bladder (2), Cervical (1), Mature B-Cell Neoplasm (1), Prostate (1) |
| H2BD (*H2BC5*) | D51/H/N/Y<br>D68H/N/G<br>E76K/Q | Prostate (3), Esophagogastric (1), Small Cell Lung (1), Soft Tissue Sarcoma (1)<br>Bladder (1), Colorectal (1), Head and Neck (1), Non-Small Cell Lung (1)<br>Bladder (4), Breast (2), Non-Small Cell Lung (2) |
| H2BN (*H2BC15*) | S55L<br>M62I/T | Bladder (1), Breast (1), Head and Neck (1)<br>B-Lymphoblastic Leukemia (1), Colorectal (1), Endometrial (1) |
| H2BC (*H2BC6*) | P10S/Q<br>H49L<br>A77T/V | Melanoma (3), Endometrial (1)<br>Non-Small Cell Lung (3)<br>Bladder (1), B-Lymphoblastic Leukemia (1), Colorectal (1) |
| H2BC (*H2BC7*) | E76K | Head and Neck (2), Bladder (1) |
| H2BH (*H2BC9*) | D68N/H<br>E76K/Q<br>E113K/Q<br>S123C | Bladder (2), Breast (1), Endometrial (1), Non-Small Cell Lung (1)<br>Non-Small Cell Lung (3), Breast (2), Bladder (1), Endometrial (1)<br>Breast (1), Cervical (1), Colorectal (1), Head and Neck (1), Non-Small Cell Lung (1)<br>Non-Small Cell Lung (6) |
| H2B2E (*H2BC21*) | F70L | Bladder (4), Breast (3), Head and Neck (2), Cervical (2), Anal Squamous (1) |

### H2B variants alter nucleosome structure and DNA accessibility

Histone variants modulate chromatin structure by influencing DNA-histone or histone-histone contacts [48]. When we consider H2B variant amino acid sequences (**Fig 1A**) and overlay these changes on a solved nucleosome structure (PDB 2CV5) [48] (**Suppl Fig 6A**), it is reasonable to suspect that the V40I substitution in H2BB, H2B2E, H2BJ, and H2BO might alter histone-DNA interactions, while the S125A substitution in H2BJ and H2BK might affect histone-histone interactions (specifically with H2AR21). Nucleosomes also breathe, and transiently expose DNA sites at each end of the nucleosome [49–52]. We therefore used several methods to understand how H2B variants impact nucleosome structure and DNA accessibility.

We first used an established, paired Cy3-Cy5 mononuleosome FRET system ([50] and **Suppl Fig 6B**) to identify how H2B variants influence nucleosomal DNA partial unwrapping. We labeled the 601 Widom nucleosome positioning sequence with the donor Cy3 fluorophore, and attached a Cy5 acceptor fluorophore to H2A (K119C) (**Suppl Fig 6B**). Because FRET efficiency is sensitive to the distance between the Cy3 label on the DNA and the Cy5 label on the histone octamer, changes in FRET efficiency can be attributed to the wrapping/unwrapping of the DNA from the octamer. Next, we reconstituted five different nucleosomes that contained H2BC (canonical H2B), H2BB, H2BD, H2BJ, or H2BK. These H2B variants were dysregulated during iAs-transformed BEAS-2B cells (from [53]), and represent all the amino acid changes found in H2B. We confirmed successful labeling and nucleosome reconstitutions by SDS and native PAGE (**Suppl Fig 6C-E)**, and then investigated nucleosome stability with NaCl titrations (0 -1.75 M). We found that the NaCl concentration that reduced FRET efficiency by 2-fold was similar to all H2B variants except for H2BJ, which required slightly more NaCl (**Suppl Fig 6F**). This suggests that H2BJ enhances nucleosome stability by wrapping DNA more tightly than in the other nucleosomes, and possibly “primed” them for reduced accessibility to transcription factors.

To test this hypothesis, we incorporated a Gal4 binding site at positions 8-26 in the 601 nucleosome positioning sequence (as described in [54, 55]; **Fig 3A**), and determined the FRET efficiency for a Gal4-DNA binding domain (Gal4-DBD) titration (the first 147 amino acids of the Gal4 transcription factor). In this measurement, Gal4-DBD binds to partially unwrapped nucleosomes, which traps them in a low FRET efficiency state. The FRET efficiency for the Gal4-DBD titration is fit to a binding isotherm, which is then used to determine the Gal4-DBD concentration that reduces FRET efficiency by 50% (*S_1/2_*). H2B variants that reduce DNA accessibility will alter Gal4 binding equilibrium to its cognate binding site, which will result in an increase in *S_1/2_.* Gal4-DBD titrations (0 -100 nM) were carried out separately with 1 nM of each nucleosome preparation at a normal physiological salt concentration. Nucleosomes containing H2BK and H2BJ had significantly increased *S_1/2_* values relative to canonical (H2BC) nucleosomes (**Fig 3B-G**; 1.6-and 3.5-fold differences, respectively), indicating that H2BK and H2BJ variants decrease DNA accessibility to Gal4. Interestingly, H2BK differs from canonical H2BC by only one amino acid substitution (S125A), while H2BJ has two amino acid substitutions (V40I and S125A; **Fig 1A**), yet the *S_1/2_* for H2BJ nucleosomes was ∼2-fold higher than for H2BK nucleosomes. We thought that the H2BJ I40 residue may contribute to the increased *S_1/2_* in HB2J relative to HB2K, so we carried out FRET efficiency measurements of Gal4-DBD with H2B2E (which only contains the V40I substitution but a canonical S125 residue). Surprisingly, the H2B2E *S_1/2_* was similar to the canonical H2BC *S_1/2_* (**Fig 3F-G**). These data imply that that the two amino acid substitutions do not function independently to decrease DNA accessibility to Gal4-DBD binding within partially unwrapped nucleosomes. These results are also consistent with our observation that amino acid substitutions in H2BB (which include V40I but not the S125A) do not impact DNA accessibility to Gal4-DBD. Instead, while an S125A substitution alone modestly reduces DNA accessibility, the impact of the V40I substitution requires S125A to generate the full 3.5-fold reduction in DNA accessibility. Thus, V40I and S125A function cooperatively to reduce DNA accessibility, most likely through reduced unwrapping.

**Fig 3.**
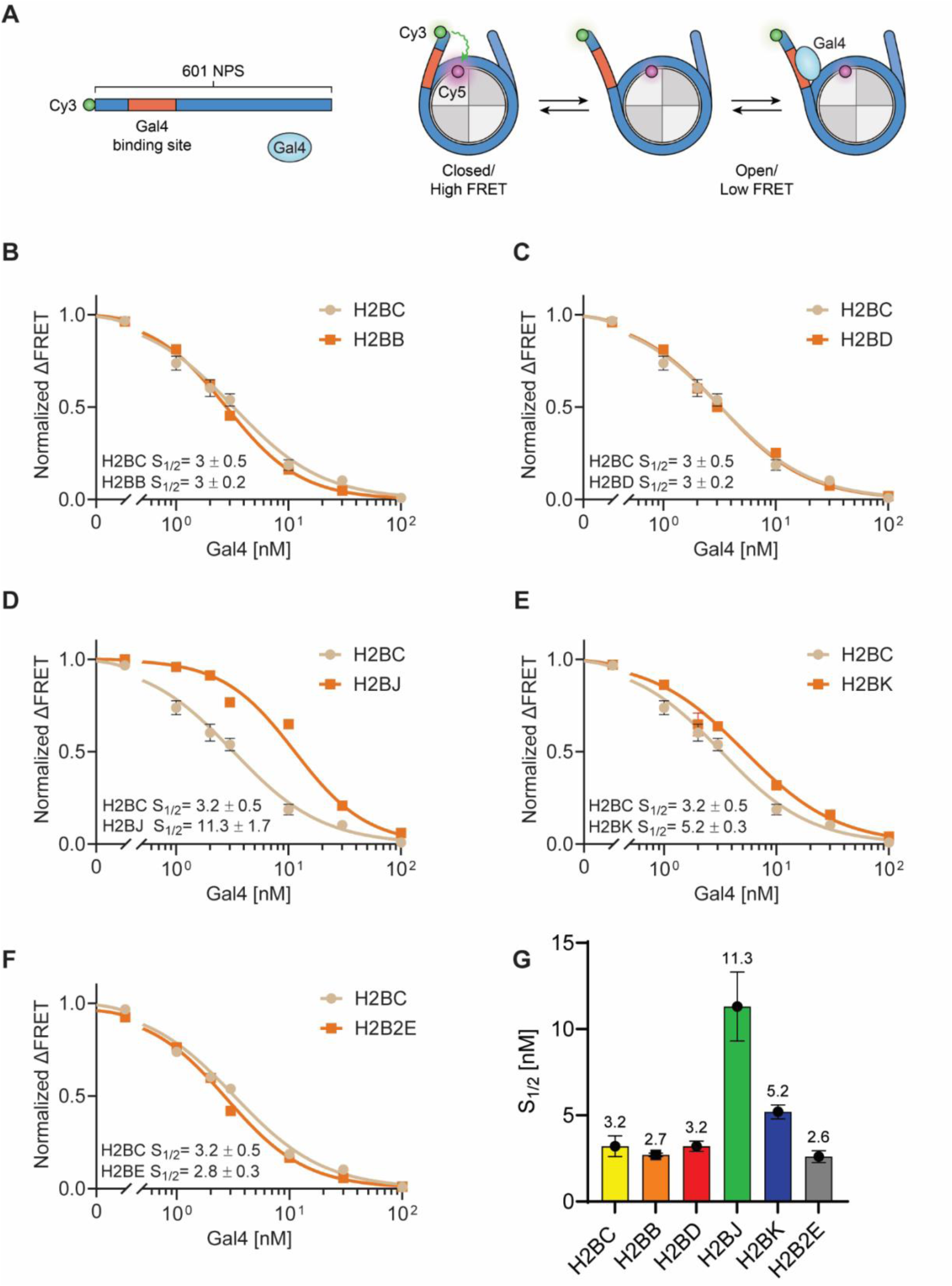
Histone H2B variants alter transcription factor access to nucleosome DNA. **A**) Schematic representation of i*n vitro* nucleosome reconstitution with Gal4, Cy3-labeled DNA, and Cy5-labeled H2A. **B-F**) Normalized ensemble FRET measurements of Gal4 binding to its corresponding binding site (position 8-26 on the DNA). Each experiment was repeated three times. **G**) Calculated *S_1/2_* values for the plots in **B-F**. Error bars indicate +/- SEM.

Because H2BJ and H2BK nucleosomes significantly altered Gal4 access to nucleosomal DNA, we asked if they would also modulate access to linker DNA, or change the trajectory of the DNA as it enters and exits the nucleosome [56]. To address this question, we reconstituted H2B variant nucleosomes whereby the 601 nucleosome positioning sequence contained an α ^32^P-label and 50 bp linker on each end (**Fig 4A**; **Suppl Fig 7A, B**). We then performed a time-dependent digestion of each nucleosome preparation (400 ng) with fixed micrococcal nuclease (MNase) concentration, and detected digestion products by native PAGE. DNA in H2BC (canonical H2B), H2BB, H2BD and H2B2E nucleosomes was readily digested to the 147 bp nucleosome core within 30 sec (**Fig 4B-E**; **Suppl Fig 7C-I**). On the other hand, it took up to 8 min to completely digest the DNA in H2BJ and H2BK nucleosomes (**Fig 4B-E**; **Suppl Fig 7G-I**). Purification, cloning, and sequencing of the intermediate digestion products indicated that the initial MNase cuts (∼ 220 bp fragment) are fairly symmetrical on each side of the nucleosome, but that subsequent cuts (past the 147 bp core) are asymmetrical (**Suppl Fig 7J**). These data are consistent with previous studies showing that nucleosomes exhibit asymmetrical breathing [57–59]. Together, these data suggest that amino acid changes found in certain H2B variants alter Gal4-DBD access to the DNA, and possibly change the trajectory of the DNA that exits the nucleosome. They also suggest that histone H2B variants could alter higher-order chromatin organization.

**Fig 4.**
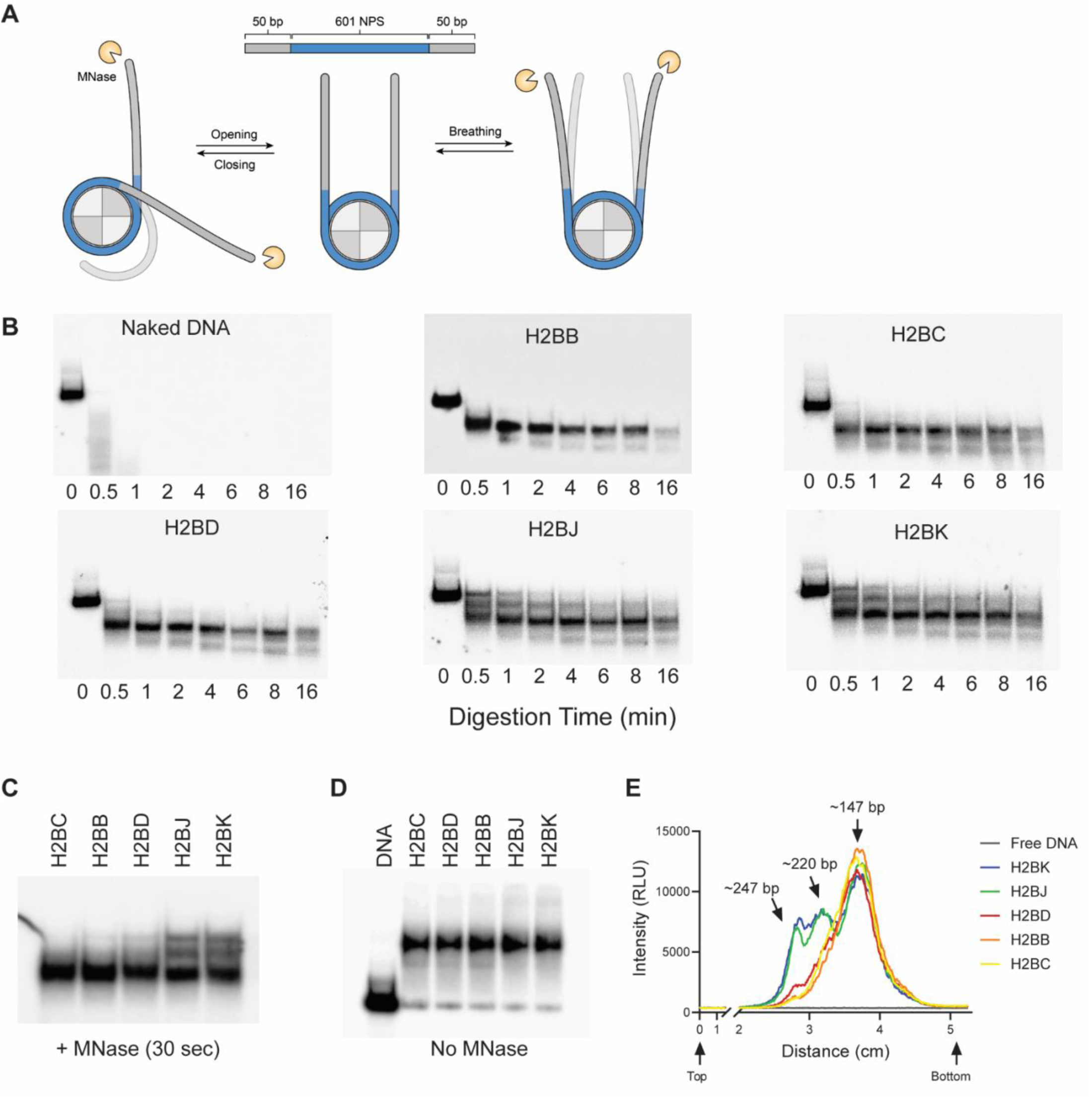
H2B variants alter access to nucleosome linker DNA. **A**) *In vitro* nucleosome reconstitution with ^32^P-labeled DNA, showing how altered DNA trajectory could block access to the linker DNA. MNase only digests accessible/naked DNA. **B**) Native PAGE analysis of MNase-digested DNA in H2B variant nucleosomes. The H2B variant is indicated in each panel. Each experiment was replicated 3 times. Grayscale images were inverted and contrast adjusted to make digestion products more visible. **C, D**) Native PAGE for H2B variant nucleosomes treated for 30 sec with (**C**) or without (**D**) MNase digestion. **E)** Densitometry scans for digested products from **C**, showing primary digestion products at ∼147 bp, ∼220 bp, and ∼247 bp. RLU = relative light units. “Top” and “bottom” are relative to the gel image in **C**. The intensity of each band was normalized to the 247 bp fragment at t = 0. Grayscale images gel images were inverted and contrast-adjusted to make digestion products more visible.

### H2B variants can alter higher-order chromatin structure

The eukaryotic genome is organized within a higher-order structure of repeating nucleosomes connected by linker DNA. To understand how H2B variants might influence this higher-order structure, we created 3kb chromatin “arrays” consisting of identical H2B variant octamers with 17 repeats of the 601-nucleosome positioning sequence, where each nucleosome was separated by a 30 bp linker [60]. We then probed the physical properties (height, Ferret’s diameter, and volume) of each variant nucleosome array by single molecule atomic force microscopy (AFM) [61]. Each variant nucleosome formed organized nucleosome and chromatin array clusters (**Suppl Fig 8)**. We then measured individual nucleosomes within these arrays, and found that H2BB, H2BD, H2BK, and H2B2E nucleosomes were significantly shorter (height) and smaller/more compact (diameter and volume) than canonical H2BC nucleosomes, while H2BJ nucleosomes were significantly slimmer (diameter) than the canonical nucleosomes (**Fig 5A-C**). When we measured the arrays, H2BB and H2BD arrays were significantly shorter than the other arrays, while H2BB, H2BJ, and H2BK arrays were significantly broader (diameter; **Fig 5D-E**). These data are consistent with the FRET and MNase data showing that H2B variants alter the structure of individual nucleosomes, and suggest that a more compact chromatin could contain more of these (more compact) nucleosomes than canonical H2BC chromatin. We might therefore expect that these differences in chromatin organization and compaction could influence chromatin function and gene accessibility.

**Fig 5.**
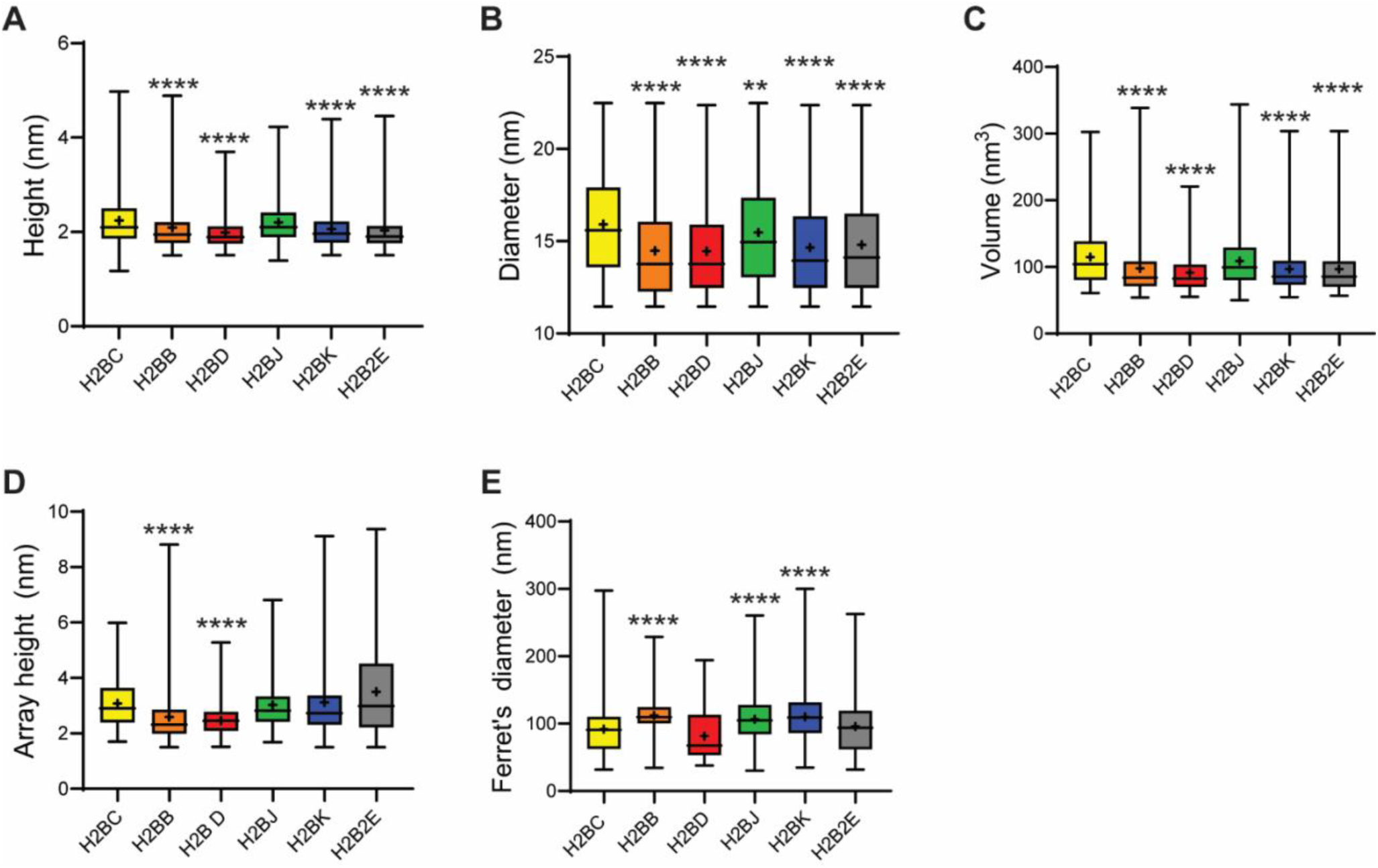
H2B variants alter higher order chromatin structure. Freshly reconstituted 17-mer nucleosome repeats (chromatin arrays) were deposited on freshly cleaved and functionalized mica, and then imaged by atomic force microscopy (AFM) images in non-contact tapping mode (in air). Nucleosome height (**A**), diameter (**B**), volume (**C**), and array (**D**) height and (**E**) diameter were determined using ImageJ software. Each experiment was repeated 3 times. **p<0.01, ****p<0.0001, as determined by Welch’s unequal variances t-test. Example AFM images are in **Suppl** Fig 8.

### H2B variants alter chromatin accessibility *in vitro*

The experiments above include nucleosomes with identical copies of each H2B variant, and canonical H2A, H3, and H4. However, histone variant combinations or heterotypic nucleosomes likely exist in cells. To study the effects of H2B variants in a physiological context, we generated a minigene containing 2xHA-tagged H2B variants under their own promoter (**Fig 6A**), and transfected these into BEAS-2B cells. We also transfected cells with a minigene containing one of the H2BC genes, so that canonical H2BC would be naturally replaced by H2B variants during cell division and growth. We maintained cells in selective medium for 20 days before measuring their expression and incorporation (**Fig 6B-D**; **Suppl Fig 9**).

**Fig 6.**
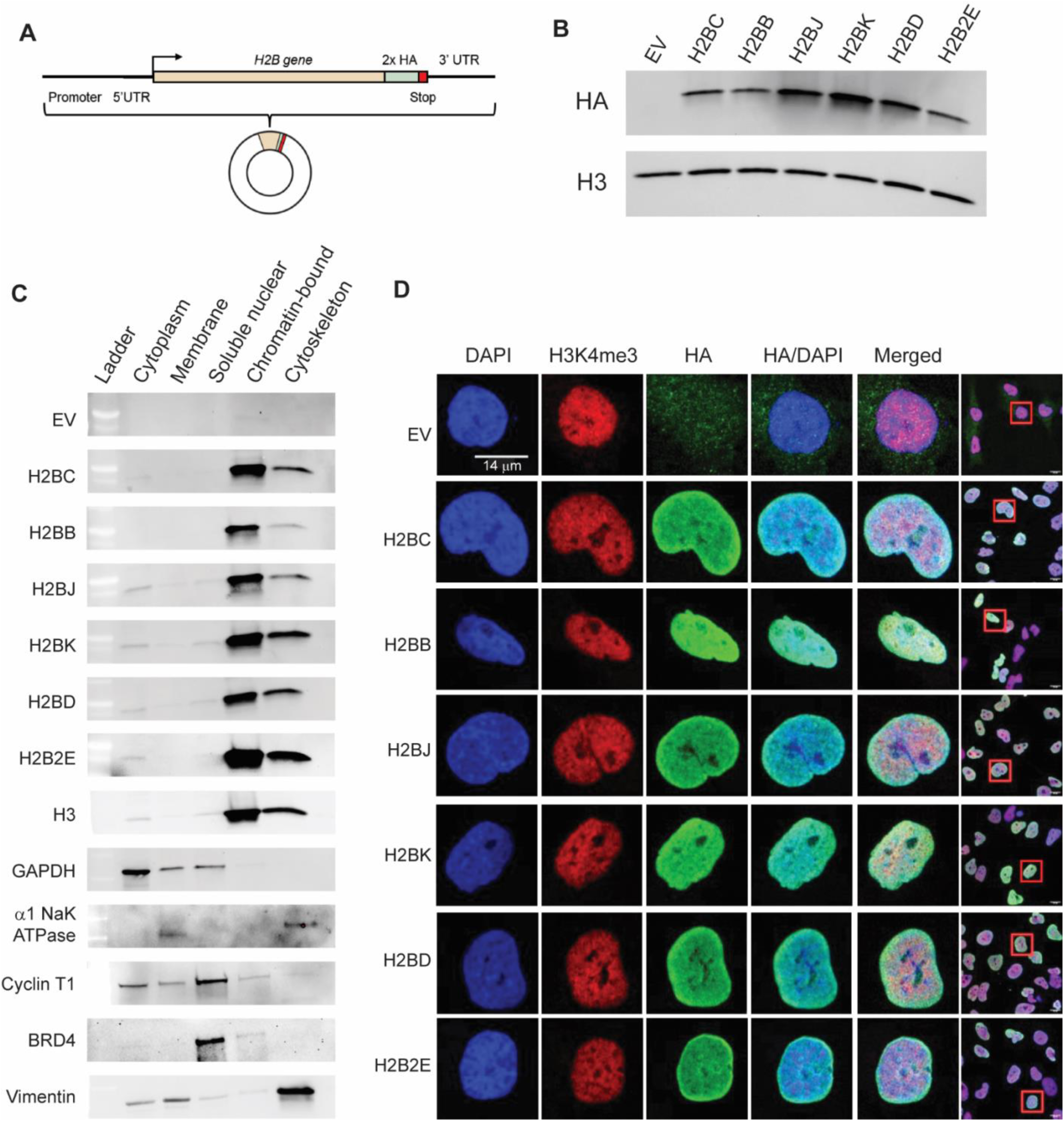
H2B variants incorporate into chromatin. **A**) Generic structure of H2B variant “minigenes” that were transfected into BEAS-2B cells, and expressed *in vitro.* **B**) Representative Western blot showing *in vitro* expression of HA-tagged H2B variants in BEAS-2B cells. H3 expression served as an internal, positive control. **B)** Representative Western blots showing that HA-tagged H2B variants are naturally incorporated into chromatin. Cytoplasmic, membrane, soluble nuclear, chromatin-bound nuclear, and cytoskeletal extracts were collected. Known localization positive controls were: GAPDH = cytoplasm; α1 NaK ATPase = membrane; Cyclin T1 and BRD4 = soluble nuclear; H3 = chromatin; Vimentin = cytoskeleton. **D**) Confocal microscopy images of individual BEAS-2B nuclei (enlarged from the cells circumscribed by red squares in the far right column), showing H2B variant expression within the nucleus.

We then performed ATAC-seq to investigate how H2B variants alter chromatin accessibility at regulatory elements. Incorporating H2B variants into chromatin altered global chromatin accessibility, where cells containing H2BD and H2B2E generated the most differentially accessible regions (DARs) and clustered together (**Fig 7A-B**; **Table 2**; **Suppl Fig 10A-B**). This result supports a recent study showing that H2BE specifically increases chromatin accessibility in the context of synaptic gene regulation and long term-memory [62]. Generally, open chromatin predominantly occurred at intronic and distal intergenic regions for most H2B variants, while H2BC open chromatin was predominantly at promoters (**Fig 7C**, left panel). Conversely, closed chromatin was primarily at promoter regions (except for H2BB and H2BC; **Fig 7C**, right panel). We posited that these intergenic regions of open chromatin may serve as enhancers, so we mapped ATAC peaks to ENCODE candidate cis-regulatory elements (cCREs) [63]. We found that most of the open chromatin regions for all H2B variants occurred at distal enhancer like signatures (dELS), while H2BC DARs were at promoter like signatures (pLS) in open chromatin and at dELS in closed chromatin regions (**Fig 7D-F**). Gene set enrichment analysis (GSEA) revealed that DARs at gene promoter regions are enriched for Yap signaling, EMT, E2F targets, TGFB, and chromatin remodeling pathways and factors (**Fig 7G-H**; **Suppl Fig 10C-D**). Given the observed accessibility changes induced by H2B variants, we predict that they lead to differential gene expression.

**Fig 7.**
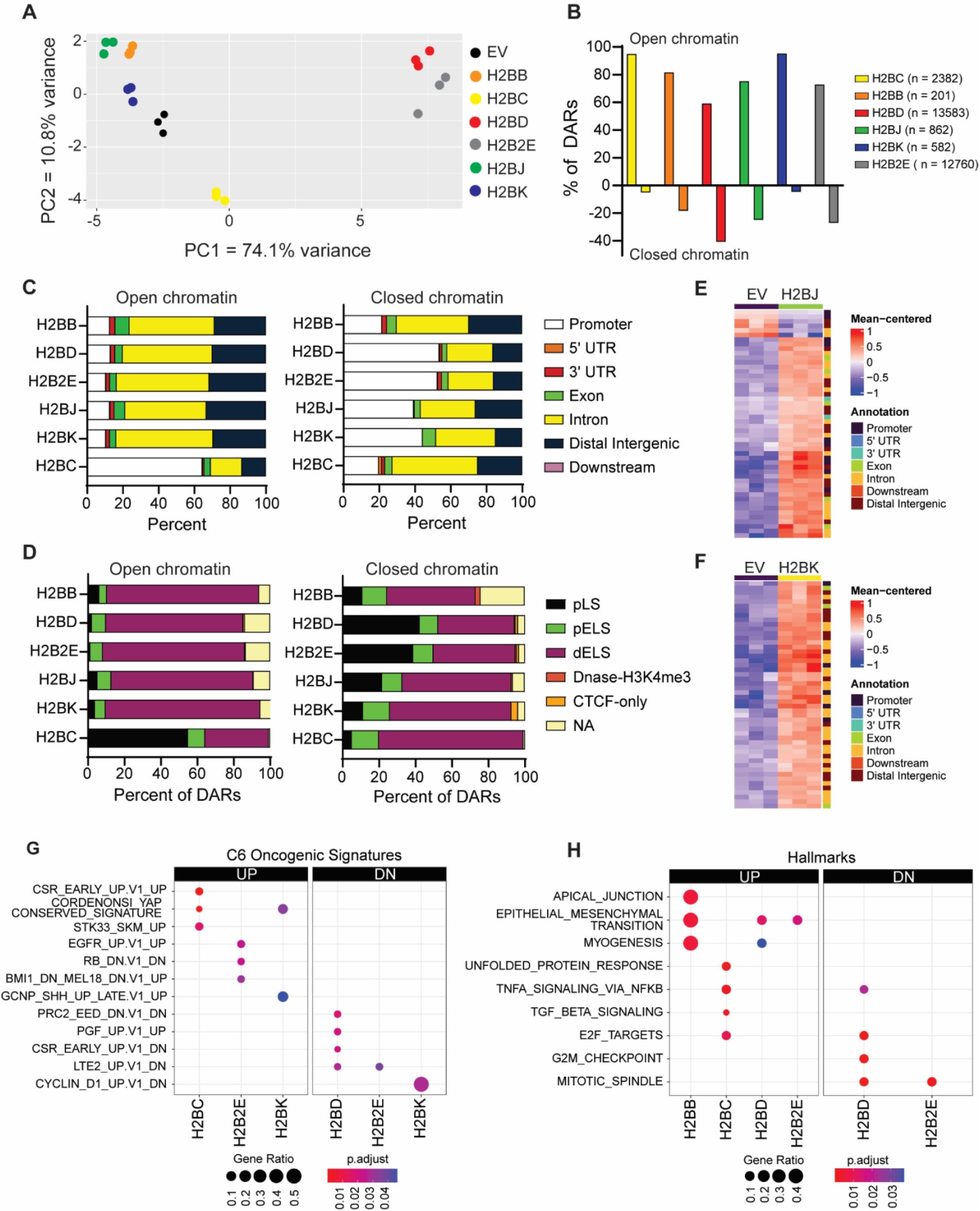
H2B variants alter chromatin accessibility. **A**) PCA plot of differentially accessible regions (DARs) in BEAS-2B containing H2B variant nucleosomes. **B)** Proportion of opened vs closed DARs in BEAS-2B cells containing H2B variant nucleosomes. DARs were defined by FDR <0.1 and Shrunken Log_2_(FC) > 1 for opened DARs, and Shrunken Log_2_FC < -1 for closed DARs. The total number of DARs is indicated in the legend parentheses. Distribution of DARs based on (**C**) genic features and **D)** ENCODE candidate cis-regulatory elements (cCREs). pLS = promoter-like signatures; pELS = proximal enhancer-like signatures; dELS = distal enhancer-like sequences. **E)** Heatmap and genic distribution of the top 50 DARs in BEAS-2B cells expressing H2BJ vs those cells containing an empty vector. **F)** Same as panel **E**, except for cells expressing H2BK. **G, H)** Dotplot of top enriched (**G**) MsigDB C6 oncogenic signatures and (**H**) hallmarks in opened (UP) or closed (DN) DARs in BEAS-2B cells containing H2B variant nucleosomes.

**Table 2.** DARs induced by histone H2B variants in genic regions.

|  | **H2BC** | **H2BK** | **H2BJ** | **H2B2E** | **H2BD** | **H2BB** |
| --- | --- | --- | --- | --- | --- | --- |
| **Open Chromatin** | 2261 | 555 | 648 | 9305 | 8039 | 164 |
| **Closed Chromatin** | 121 | 27 | 214 | 3455 | 5544 | 37 |
| **Total** | 2382 | 582 | 862 | 12760 | 13583 | 201 |

### H2B variants alter oncogenic gene expression programs in lung epithelial cells

We next performed RNA-seq to quantify gene expression in each of the transfected cell lines generated above. We observed that only H2BB (*H2BC3*) was over-expressed, most likely because of the low basal *H2BC3* expression (**Suppl Fig 11A**). Importantly, incorporating H2B variants into chromatin significantly altered gene expression, with gene expression profiles from H2BD and H2B2E grouping together and apart from the rest (**Fig 8A**, **Suppl Fig 11B-C**) but nevertheless sharing many differentially expressed genes with cells incorporating other H2B variants (**Fig 8B**). Gene set enrichment analysis (GSEA) revealed activating and suppressing terms related to tumorigenesis (e.g., MYC, KRAS, TNFA, TGFB), cell proliferation, and EMT signatures (**Fig 8C-D**; **Suppl Fig 11D-E**). Even with these activated or suppressed oncogenic pathways, however, we did not detect any significant differences in cell migration or proliferation (**Suppl Fig 12A-C**). This result was somewhat surprising, and suggests that oncogenic transformation (at least in BEAS-2B cells) requires something more than an activated or suppressed oncogenic pathway.

**Fig 8.**
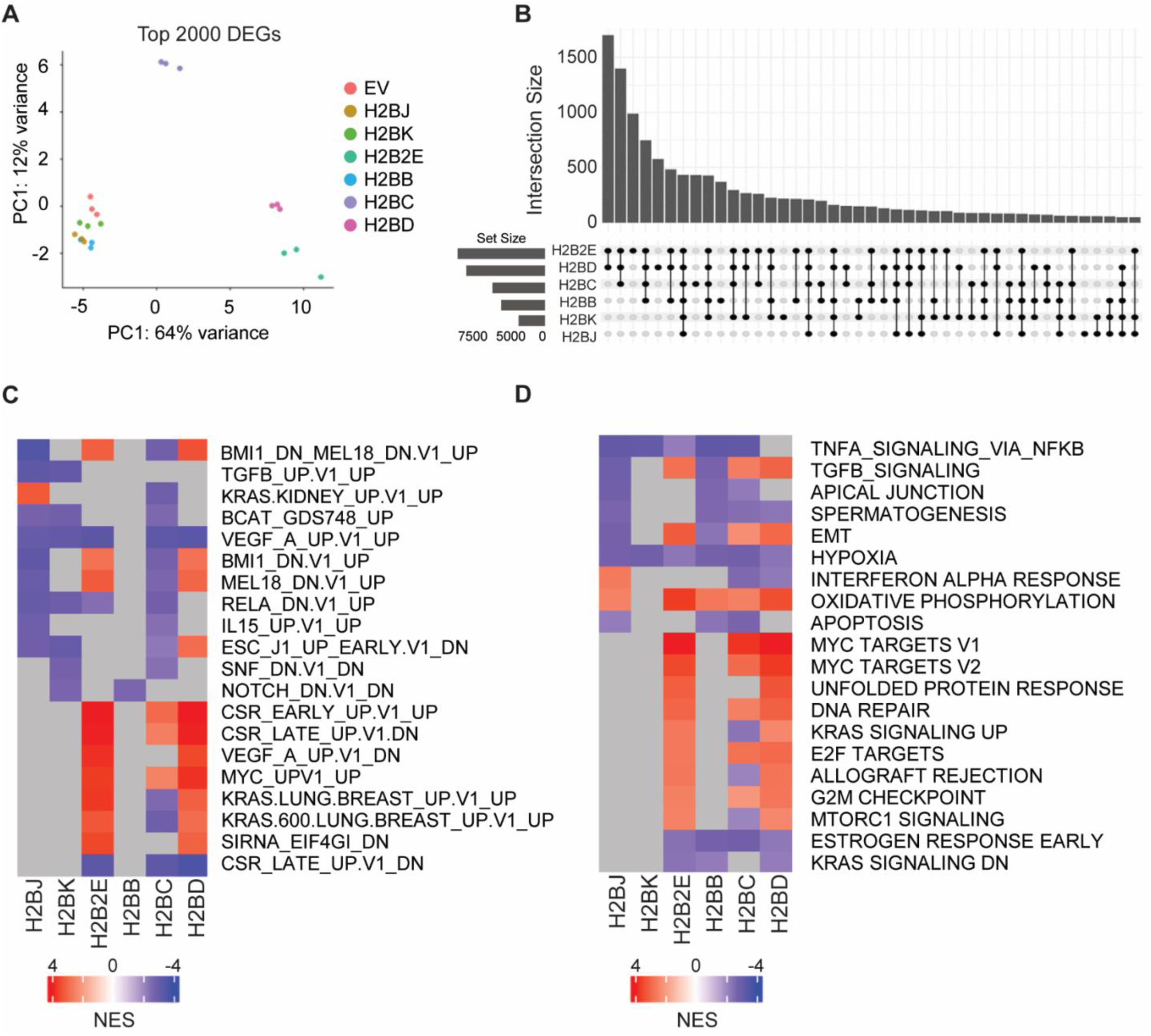
H2B variants alter oncogenic pathways in lung epithelial cells. .**A**) PCA plot of BEAS-2B gene expression patterns after incorporating H2B variants into nucleosomes/chromatin. **B**) Upset plot of unique and overlapping DEGs in BEAS-2B cells containing H2B variant nucleosomes. Black dots and connecting lines indicate shared associations. Set size = number of DEGs in each overlapping set, intersection size = number of DEGs shared in that intersection. **C, D)** Heatmaps of the top 20 enriched (**C**) MsigDP oncogenic signatures and (**D**) hallmarks for H2B variant DEGs. NES = normalized enrichment score. Grey = no hits for the associated pathway.

We then merged ATAC-seq data, and annotated DARs to nearby genes and their corresponding RNA-seq expression levels. For some H2B variants, there was substantial overlap between open/closed chromatin and up/down-regulated gene expression, while others did not show any overlap (**Suppl Fig 13A**). For example, H2BJ expression increased chromatin accessibility at *ITGAX* and decreased it at *FCRLA* promoters, corresponding to increased and decreased gene expression, respectively (**Fig 9A**). H2BK expression showed the same pattern at the *PLK2* (increased) and *KRT17* (decreased) promoters (**Fig 9B**). For other variants, there was little to no association between DARs and gene expression, which might imply that these loci are regulated by distal elements and post-transcriptional regulatory mechanisms [64, 65]. Where chromatin accessibility did correlate with gene expression, these genes were enriched for Yap signaling, E2F targets, and chromatin remodeling factors, consistent with our above findings (**Fig 9C-D**; **Suppl Fig 13B-C**).

**Fig 9.**
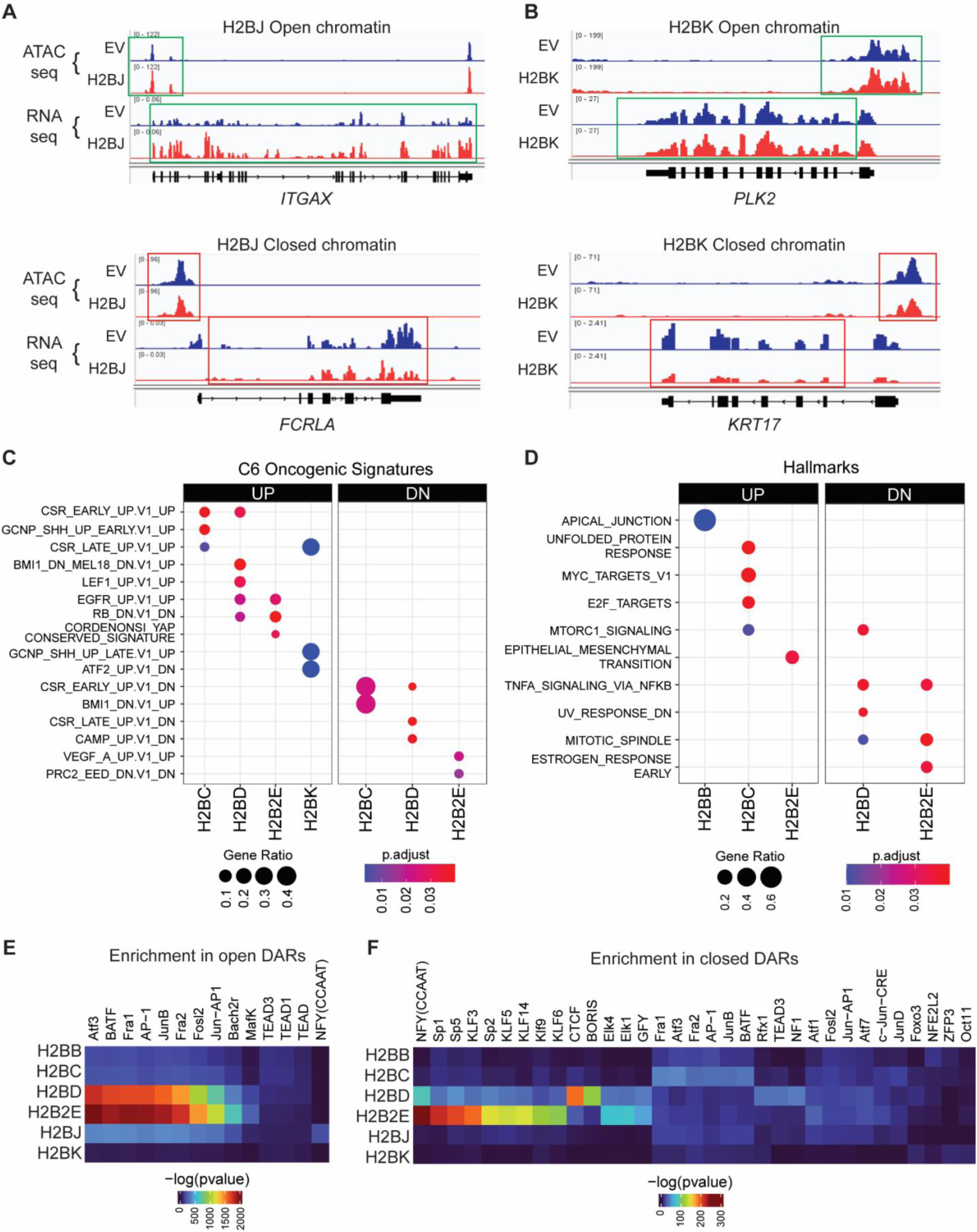
H2B variants alter chromatin accessibility and gene expression. Genome browser tracks for (**A**) H2BJ and (**B**) H2BK DARs where an opened DAR was associated with increased gene expression (top panels), and a closed DAR with decreased gene expression (bottom panels). **C, D)** Dotplot of top enriched (**C**) MsigDB C6 oncogenic signatures and (**D**) hallmarks where opened (UP) and closed (DN) DARs correspond to increased and decreased RNA expression of at least five overlapping genes, respectively. Genes with differentially accessible promoters were compared with differentially expressed genes at a cutoff of 0.05 adjusted p-value. Transcription factor motif enrichment of these overlapping DEGs in opened (**E**) and closed (**F**) DARs.

We also performed motif enrichment analysis on the DARs, and found significant enrichment for the bZip family of transcription factors in opened DARs, including AP-1 and oncogenic FOS, JUN, ATF, and MAFP transcription factors [66–69], especially in cells expressing H2BD and H2B2E (**Fig 9E**). In contrast, motifs for oncogenic Kruppel-like transcription factors and specificity protein (KLF/SP) family members (e.g., KLF1, KLF3, KLF5, SP1, and SP2) [70–73] were enriched in closed chromatin (**Fig 9F**). We also observed enrichment of the CCCTC-binding factor (CTCF) motif in closed chromatin regions, a factor involved in maintaining chromatin structure and regulating gene expression [74, 75]. These findings highlight the differential role of H2B variants in modulating chromatin accessibility, and suggest a profound impact on transcriptional regulation and potential oncogenic pathways.

## DISCUSSION

Alterations to chromatin structure can arise from histone gene mutations, post-translational modifications, or incorporation of histone variants, all of which are associated with cancer initiation and progression [10]. In our prior work, we found that H2B variants were up-regulated during iAs-induced oncogenic transformation [53]. Here, through a pan-cancer analysis, we found that H2B variant dysregulation was associated with many cancers and 5-year prognoses (**Fig 1**), similar to what has been found with H2A and H3 variants [19, 21, 76–79]. We also found that H2B variants had distinct mutational hotspots (**Fig 2**), some of which might affect DNA-histone or histone-histone interactions [43–45, 47]. These findings prompted us to determine how H2B variants influence chromatin architecture, dynamics, and gene expression programs.

Our use of FRET experiments, Gal4 transcription factor binding assays, MNase digestion assays, and single-molecule AFM provided a multi-faceted approach to probe the physical and functional consequences of H2B variant incorporation into chromatin. Our FRET experiments (like those in [50, 80]) suggested that most H2B variant nucleosomes breathe like canonical nucleosomes, but bulk nucleosome measurements might mask subtle differences in nucleosome dynamics. We therefore repeated the experiments with a Gal4-DBD transcription factor binding assay, and found that H2BJ and H2BK nucleosomes significantly reduced Gal4-DBD access to nucleosomal DNA where the amino acid substitutions V40I and S125A function cooperatively. Nucleosome breathing (Gal4-DBD accessibility) results were corroborated by the MNase experiments, where linker DNA in H2BJ and H2BK nucleosomes was more “protected” from enzymatic digestion than DNA in the other nucleosomes. These data are consistent with other studies showing that histone variants may change the trajectory of linker DNA, and its interactions with the nucleosome core [56]. In some H2B variant nucleosomes, we also saw evidence for asymmetric accessibility to linker DNA, suggesting that “under tension” one side of the nucleosome is more flexible [57, 81, 82]. Unfortunately, the MNase experiments do not identify which “end” of the nucleosome is more or less protected, nor are they precise enough to detect asymmetric accessibility differences between the H2B variant nucleosomes. Finally, single molecule analysis of nucleosomes using AFM showed that several H2B variant nucleosomes (and chromatin) were more compact than canonical nucleosomes and chromatin.

We speculated that the physical differences between H2B variant nucleosomes would be reflected as DARs and differentially expressed genes (DEGs) *in vitro*. Others have used specific cancer cell models and antibodies to correlate histone variants with gene expression [83]. Unfortunately, there are no commercial antibodies that are specific for each H2B variant, so we instead used a minigene expression system to incorporate H2B variants into chromatin, making sure to avoid over-expressing each histone (which might otherwise cause DNA damage or chromosome instability [84, 85]). Incorporating H2B variants into chromatin altered chromatin accessibility at cis-regulatory elements, and at genes involved in chromatin organization, DNA repair, and oncogenesis. Curiously, BEAS-2B cells expressing H2B variants and dysregulated oncogenic pathways did not show corresponding oncogenic phenotypes (i.e., increased cell migration or proliferation) as might be expected (e.g., [35]). Thus, one possibility is that the H2B variants studied here do not act as “oncohistones” *per se*, or perhaps the histone variant abundance in this artificial system may not be high enough for BEAS-2B cells to exhibit oncogenic properties. Based on our data, however, we propose an alternative hypothesis that H2B variants may influence early-stage, cancer-associated regulatory mechanisms that "prime" cells for potential transformation by some other trigger (e.g., iAs). Consistent with this notion, others have shown that certain genetic or epigenetic alterations can initiate oncogenic pathways without immediately transforming cells, but rather change chromatin dynamics and gene expression in ways that predispose cells to future oncogenic changes [86–91].

In summary, this study is the first to systematically analyze how histone H2B variants alter nucleosome structure and gene expression/chromatin accessibility in cells. We show that H2B variants shape chromatin architecture and gene expression, and that naturally occurring histone variants and epigenetic mechanisms regulate oncogenic pathways even in the absence of any DNA mutation or external insult. While our findings suggest that H2B variants might "prime" cells for cancer, this remains a hypothesis requiring further validation. Nonetheless, our study represents the first interdisciplinary effort aimed at understanding the biochemistry and functional outcomes of H2B variants.

## STAR★Methods

### Key resources Table

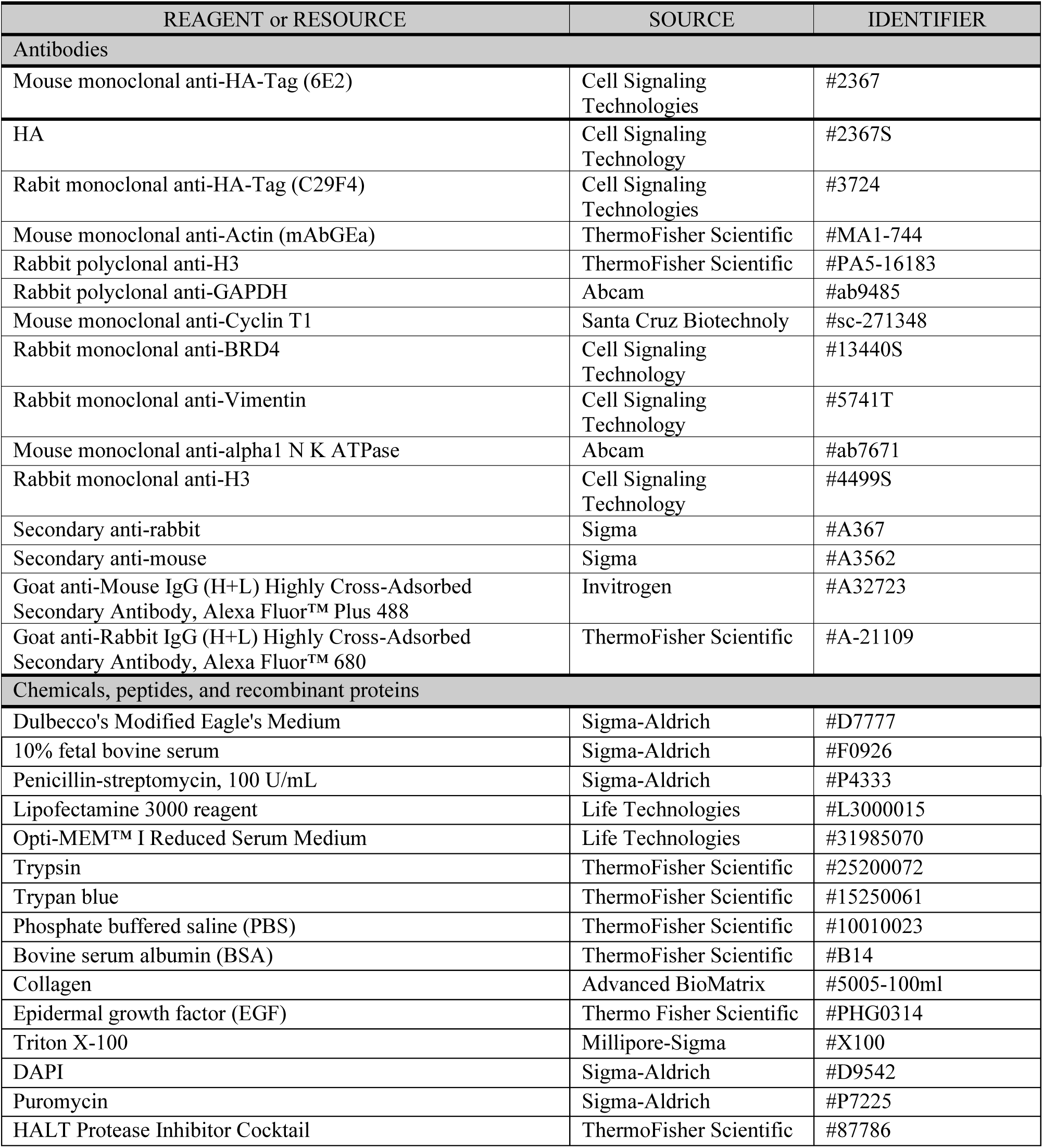
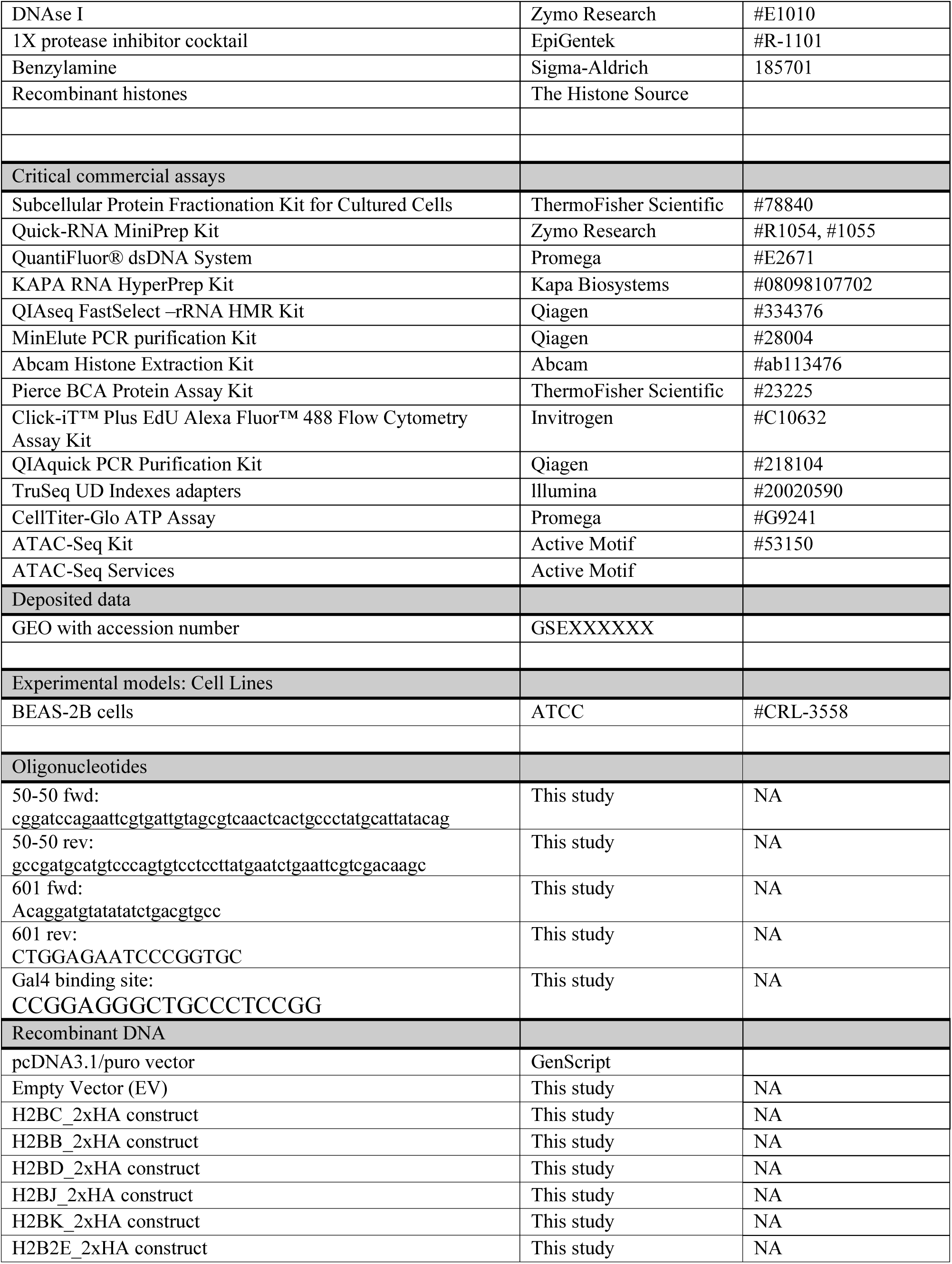
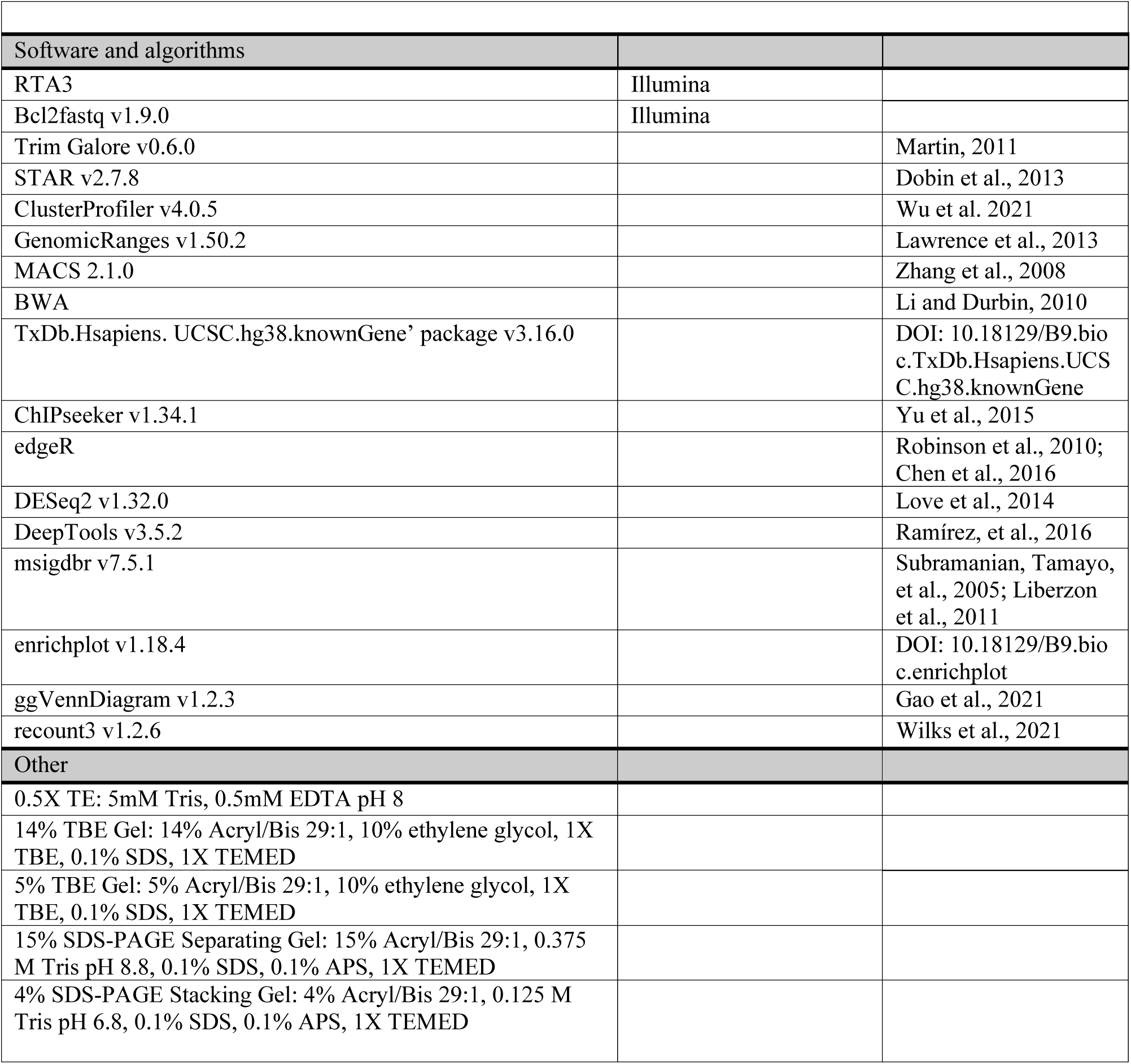

## RESOURCE AVAILIBILITY

### Lead contact

Further information and requests for resources and reagents should be directed to and will be fulfilled by the lead contact, Yvonne Fondufe-Mittendorf.

### Materials availability

This study did not generate new unique reagents.

### Data and code availability

- RNA-Seq and ATAC-Seq data pertaining to this paper have been deposited at GEO and are publicly available as of the date of publication. The accession number is listed in the key resources table.
- This paper does not report any original code.
- Any additional information required for the re-analysis of the data reported here can be requested from the lead contact.

### Experimental model and study participant details

N/A

### Method Details

#### TCGA data processing and analyses

Read counts for TCGA samples were based on GENCODE v29 gene annotations, and were obtained using the recount3 v1.2.6 R package [92]. Per-gene raw counts and transcripts-per-million (TPM) values were calculated using the ‘compute_read_counts’ and ‘getTPM’ functions, respectively. Tumor purity estimates were obtained from the ‘TCGA_mastercalls.abs_tables_JSedit.fixed.txt’ file from ‘https://gdc.cancer.gov/about-data/publications/pancanatlas’, and merged with the count data within a single RangedSummarizedExperiment object. Only samples where tumor purity was >0.7 and the library size was > 15 million reads were retained. Samples with the same sample ID were deduplicated by retaining only the sample with the highest number of uniquely mapping reads. To simplify interpretation of the results and to prevent duplicated patients, samples labelled as "Additional - New Primary", "Recurrent Tumor", or "Metastatic" were removed. To remove lowly expressed genes, only genes with >15 read counts in >16 samples (the sample size of the least represented cancer type) were retained. Variance stabilizing transformed (VST) counts were obtained using the ‘vst’ function in DESeq2 v1.32.0, with the parameter ‘blind=FALSE’ and the design set to ‘∼Cancer_Type’ [93].

For a given cancer type, only genes with ≥10 read counts in at least half of the samples were analyzed. We used the VST counts and the ‘corr.test’ function in the psych v2.2.5 package to generate Spearman correlations for each gene and H2B variant. For each cancer type, H2B variant, and gene combination, p-values were corrected for multiple testing using the Benjamini-Hochberg method (*p_adj_* ≤0.05). Genes were then ranked based on the absolute value of the correlation. For GSEA, ranked genes were used as the input to ClusterProfiler v4.0.5 [94].

To identify differentially expressed genes in adjacent normal tissue versus the corresponding tumor, we used the recount3 package to obtain per-gene read counts as described above, but the downstream preprocessing was done separately from the tumors-only analysis. Here, duplicated samples were removed by keeping only the sample with the highest number of uniquely mapped reads. Samples with <15 million uniquely mapped reads were removed, as were FFPE samples. Samples labelled as "Primary Tumor" or "Solid Tissue Normal" were retained, and only the cancer types with ≥15 samples in each group were processed. Raw counts were calculated using the ‘compute_read_counts’ function in recount3. For each cancer type, the tumor versus normal contrast was tested using the default workflow encoded in the ‘DESeq’ function with an adjusted p-value cutoff of 0.05. Gene expression data were converted to VST counts and visualized with the ‘vst’ function with the parameter ‘blind=FALSE’.

Cox proportional hazards regressions were used to estimate the association between each histone gene and the survival rate of various cancer types, and were fit via the R v4.1.0 package ‘survival’ [95, 96]. Each model was adjusted for ‘age at diagnosis’ and ‘gender’; with ‘days until last follow-up’ as the time component and ‘vital status’ (alive or dead) as the event of interest. Gene expression was modeled as log2(counts per million), and p-values adjusted using Benjamini-Hochberg method to maintain a <5% false discovery rate.

### Cell lines

Bronchial Epithelial (BEAS-2B) cells were grown in Dulbecco’s Modified Eagle Medium (DMEM) (Gibco, # 11965092) supplemented with 10% FBS (Sigma-Aldrich, #12103C) and 1% penicillin-streptomycin (Gibco, #15070063) in a 5% CO_2_ humidified chamber at 37°C, until cells reached 80-90% confluency.

### DNA Constructs

#### Fluorescence resonance energy transfer (FRET)

We used primers labeled with a Cy3 NHS ester (GE Healthcare, #GEPA13101) at a 5’ amine-modified internal thymine to PCR amplify the 147 bp nucleosome positioning sequence (NPS) fragment from a plasmid containing the Widom 601 sequence that had a Gal4 binding site (5’-CCGGAGGGCTGCCCTCCGG-3’) at bases 8-26 [54, 97, 98] (**Fig 3A**). Prior to PCR, the oligonucleotides were purified on a 218TP C18 reverse-phase column (Grace/Vydac, Avantor #218TP3215) as previously described [99, 100]. Following PCR amplification, the Cy3-labeld fragments were purified using a MonoQ column (GE Healthcare, #17-5166-01).

#### Micrococcal nuclease (MNase)

The 601 NPS (**Fig 4A**) was labeled through PCR amplification with α-^32^P (Revvity, #BLU003H250UC) using oligonucleotides which add 50 bp linker on each end (247 bp amplicon). Amplified DNA was concentrated on a 30K Amicon® Ultra-15 Centrifugal Filter Unit (Sigma, #UFC903024) at 4°C and 3000 rpm, and amplification products separated on a 14% native polyacrylamide gel in 1X TBE for 3 hours. The corresponding DNA band was cut, crushed, and incubated on a rotator in Crush-N-Soak buffer (0.75 M NH₄CH₃CO₂, 0.3 M C₂H₃NaO₂, 1mM EDTA, 0.1% SDS) at room temperature overnight. DNA was purified via butanol reduction into 0.5X TE buffer, and concentrated using a 30K Amicon® Ultra-4 Centrifugal Filter Unit (Sigma, #UFC803024). To ensure the highest purity, DNA was then dialyzed overnight at 4°C against 0.5X TE using a 10kDa Slide-A-Lyzer Mini Dialysis Unit (ThermoFisher Scientific, #0069570). DNA was then quantified using the Thermo Scientific™ NanoDrop™ OneC Microvolume UV-Vis Spectrophotometer (ThermoFisher Scientific, #840274200).

#### 17-mer repeat DNA

A plasmid containing a 17-mer of the 601 NPS was a kind gift from the Poirier lab (see [60]). Plasmid DNA was digested with ∼0.5 units/ug with DdeI (NEB, #R0175S) overnight at 37°C, and heat-killed at 65°C for 20 minutes. We checked digestion efficiency by 0.7% agarose gel electrophoresis, and then purified the digested plasmid via phenol-chloroform extraction.

### Nucleosome and chromatin assembly

#### Histone octamer preparations

Individual histone octamers were purchased from the Histone Source (Colorado State University), or refolded using individual histones as previously described [99, 101]. Briefly, lyophilized histones were resuspended in unfolding buffer (20 mM Tris-HCl pH 7.5, 7 M guanidine HCl, and 10 mM DTT) at 3-5 mg/ml for 1 hr, then centrifuged to remove aggregates. Each unfolded histone was quantified by measuring absorbance at A276 nm using a Nanodrop™ spectrophotometer. Histones (H2A:H2B:H3:H4) were combined at a ratio of 1.2:1.2:1:1 each. Samples were dialyzed in refolding buffer (10 mM Tris-HCL pH 7.5, 1 mM EDTA, 2 M NaCl, and 5 mM β-mercaptoethanol) at 4°C for a total of 3 rounds, where the first dialysis round was >12 hrs (overnight) and the others were >6 hours each. For fluorescence experiments, heterodimers were combined at a 1:1 ratio of (H2A:H2B) and (H3:H4), respectively, and refolded into octamers. All octamers were labeled with Cy5-maleimide (GE healthcare, #GEPA15130) at H2AK119C as previously described [100, 102].

As needed, heterodimers, tetramers, and octamers were purified on a Superdex 200 gel filtration column (GE Healthcare, #28990944) and AKTA FPLC instrument. Eluted fractions were analyzed by SDS-PAGE on 18 % acrylamide gels, and those fractions containing heterodimers, tetramers, or octamers were pooled and concentrated with a 30K Amicon® Ultra-4 Centrifugal Filter Unit (Sigma, #UFC803024). Samples were quantified by A276, adjusted to 50 %(v/v) glycerol, and stored at -20°C for future use.

#### Nucleosome preparation

Nucleosomes for all experiments were prepared and purified as previously described [99, 101]. Nucleosomes for FRET experiments were either purified by mixing Cy3-labeled DNA and Cy5-labeled histone octamer at a ratio of 1.25:1, or by mixing Cy3-labeled DNA, Cy5-labeled heterodimer, and tetramer at a ratio of 1.5:1.5:1. Nucleosomes for MNase experiments were purified by mixing α-^32^P labeled DNA with purified octamer at a ratio of 1.25:1. All nucleosomes were reconstituted via dialysis from a high salt reconstitution buffer (0.5X TE, 1 mM benzamidine HCl, 2M NaCl) into a low-salt reconstitution buffer (0.5X TE, 1 mM benzamidine HCl) at 4°C. After 5-6 h, the samples were placed into a new 4L low-salt buffer bucket at 4°C overnight. Dialyzed nucleosomes were purified in a 5-30% w/v sucrose gradient with a SW41 Ti rotor (Beckman Coulter, #331362) in an Optima L-90K or Optima L-100 XP ultracentrifuge (Beckman Coulter, #2043-30-1191; #8043-30-1124) at 41,000 rpm for 22h at 4°C. Samples were then fractionated and analyzed by native PAGE. Fractions containing center-positioned nucleosomes were concentrated into 0.5x TE pH 8 with a 30K Amicon® Ultra-15 Centrifugal Filter Unit (Sigma, #UFC903024) at 3000 rpm at 4°C. We verified nucleosome purity by native PAGE with a 5% acrylamide gel in 0.5x TBE. To resolve individual histones, purified mononucleosomes were also precipitated with 5 mM MgCl_2_ [103], heated at 90°C in 1× SDS loading buffer, and separated by SDS-PAGE on an 18% gel. The gel was stained with Coomassie blue, and band intensities quantified using a ChemiDoc MP Imaging System.

#### Nucleosome array preparation

Purified histone octamers and the linearized, purified, 3 kb 17-mer 601 NPS plasmid were reconstituted as nucleosome arrays by salt gradient dialysis [60]. Briefly, the linear plasmid was mixed with histone octamer at a ratio of 1:0.9 in 1x TE, 2M NaCl on ice for 30 min. The mixture was then transferred to a 3.5kDa Slide-A-Lyzer Mini Dialysis Unit (ThermoFisher Scientific, #0069558), and dialyzed at 4°C against 1X TE buffers containing 1M NaCl for 2 hours; 0.8M NaCl for 2 hours; 0.6M NaCl overnight; and 0.15M NaCl for 2 hours.

### FRET measurements

Gal4 was a kind gift from the Poirier lab, and used previously in [54, 55]. Variant-containing nucleosomes (1 nM each) were incubated for 10 min with increasing Gal4 concentrations (0-100 nM) in T130 buffer (10 mM Tris-HCl pH 8, 130 mM NaCl, 10% glycerol, and 0.25% Tween-20), and fluorescence spectra were measured with a Horiba Scientific Fluoromax 4 (λex =510 and 610 nm; λ_em_ = 530-750 nm and 630-750 nm for donor and acceptor excitations, respectively). FRET efficiency was calculated using the RatioA method [104].

### MNase digestion

Nucleosomes (n=3, 400 ng each) that were reconstituted with α-^32^P labeled DNA (above) were incubated with 0.0125 U/µl of MNase (Sigma, #N3755), with 100µl aliquots of the reaction mixture removed into 10µl of stop solution (250mM EDTA, 100mM EGTA, 0.1% SDS) at t=0, 0.5, 1, 2, 4, 6, 8, and 16 min. Digested DNA was then purified using the QIAquick PCR Purification Kit (Qiagen, #28104), and eluted in 40 µl 0.5X TE. Purified DNA was mixed with Orange G (Sigma, #O3756), and separated by native PAGE with a 5% gel in 0.5X TBE at 200V for 3 hours. Gels were placed on phosphor screen overnight, imaged with an Amersham Typhoon (GE Healthcare), and band intensities quantified using Image Quant software. Each experiment was repeated 3 times.

### Atomic force microscopy

Purified chromatin arrays (n=3) were diluted 40-fold in [35mM KCl, 15mM NaCl, 5mM HEPES and 0.25% glycerol] before being gently placed in a single layer on freshly cleaved mica that had been functionalized with aminopropyl-silantrane (APS) for 10 min and washed 5x with 1000 µl of pure water before drying with argon gas [61]. The samples were then gently rinsed 2X with 200 µl of pure water before very gently drying with argon gas. Images were obtained with an Asylum Research’s Cypher S AFM commercial atomic force microscope (Oxford Instruments) and silicon cantilevers (OTESPA-R3 from Olympus; nominal resonance of ∼300 kHz, stiffness of ∼42 N/m) in non-contact tapping mode, in air. Images were processed as previously reported [61, 105]. Image analyses were performed with Image J, and statistical analyses and visualization were done in GraphPad Prism version 10 (La Jolla, CA, USA).

### H2B minigenes and cell line transfection

H2B minigenes were created by cloning the promoter, 5’UTR, coding, HA, and 3’UTR sequences of six histone H2B genes into a promoter-less pcDNA3.1/puro vector (GenScript). Minigenes were transfected into BEAS-2B cells using 5 µg of each plasmid and lipofectamine 3000 transfection reagent (Invitrogen, #L3000015), as per the manufacturer’s protocol. After 48 hours, the transfection medium was removed and replaced with selection medium (DMEM, 10% FBS, and 1% penicillin-streptomycin supplemented with 1 ug/mL of puromycin (Gibco, #A1113802)). Cells were maintained in selective medium for 20 days by changing medium every 2-3 days, and then used for downstream experiments.

### Whole cell and histone extracts

To create whole cell protein lysates, ∼5-10x10^6^ BEAS-2B cells were resuspended in 200 µl of 1X RIPA buffer (150 mM NaCl, 1% Nonidet P-40, 0.5% sodium deoxycholate, 0.1% SDS, 25 mM Tris-HCl pH 7.4) supplemented with 1X protease inhibitor cocktail (PIC) (EpiGentek, #R-1101). Cell suspensions were intermittently sonicated for 12 cycles (30 sec on/12 sec off) in a Bioruptor*Plus* sonicator, centrifuged at 13000 rpm for 15 minutes at 4°C to remove insoluble cell debris, and the remaining supernatant (∼180 µl) transferred to fresh tubes. Histones were extracted from cell lysates using the Abcam Histone Extraction Kit (#ab113476) according to the manufacturer’s protocol. Protein concentrations in whole cell and histone extracts were determined using the Pierce BCA protein assay Kit (Thermo Fisher Scientific, #23225).

### Subcellular Fractionation

Subcellular fractions were generated with the Subcellular Protein Fractionation Kit for Cultured Cells (ThermoFisher Scientific, #78840). Briefly, ∼4x10^6^ cells were harvested and separated into cytoplasmic, membrane, soluble nuclear, chromatin-bound nuclear, and cytoskeletal fractions. Approximately 15 µl from each fraction was then separated by SDS PAGE on precast polyacrylamide gels (Bio-Rad Laboratories, 4–20% Mini-PROTEAN® TGX™ Precast Protein Gels, 10-well, 30 µl, #4561093), and transferred to nitrocellulose membranes as described above.

Western blots were then performed using an HA antibody (Cell Signaling Technology, #2367S), as described above. Positive controls for each sub-cellular fraction included: GAPDH (Abcam, #ab9485) as a cytoplasmic marker; α1-NaK ATPase (Abcam, #ab7671) as a membrane marker; Cyclin T1 (Santa Cruz Biotechnology, #sc-271348) and BRD4 (Cell Signaling Technology, #13440S) as markers for the soluble nuclear extract; histone H3 (Cell Signaling Technology, #4499S) as a marker for chromatin; and Vimentin (Cell Signaling Technology, #5741T) as a cytoskeletal marker.

### Western Blot Analyses

For western blot analyses, whole cell, histone extracts, or cellular factions were separated by SDS PAGE on precast polyacrylamide gels (Biorad, 4–20% Mini-PROTEAN® TGX™ Precast Protein Gels, 10-well, 30 µl, #4561093). Thereafter, proteins were transferred to a nitrocellulose membrane using an iBlot 2 Gel Transfer Device (Invitrogen, #IB23001). The nitrocellulose membrane was then incubated for 1 hr in 10 ml blocking buffer (1X TBST with 5% w/v nonfat dry milk). Blocking buffer was decanted, and membranes incubated with primary antibodies (in 0.5% nonfat powdered dry milk + PBST) overnight at 4°C, and then for 1 hr with secondary antibody (α-Rabbit or α-Mouse) conjugated to alkaline phosphatase. Membranes were then incubated with a fluorescent ECF substrate (Cytiva, #RPN5785), and proteins bands detected with a ChemiDoc MP Imaging System (Bio-Rad Laboratories).

### Cell migration assay

Transwell migration assays (n=3) were used to estimate cell migratory potentials. Cells were grown in complete and selective media, and changed to serum-and growth factor-free media (basal DMEM+ 0.1% BSA) before reaching confluency. After 24 hours, the starved cells were collected with trypsin, and ∼5,000 cells (in 100 µl serum free media) were seeded into the upper compartment of the transwell inserts (Greiner bio-one, # 662638) that had been treated with a coating solution (1X PBS, 0.002 mg/mL collagen, 0.042 mM NaOH) for 30-60 minutes. The lower chamber of the transwell inserts contained 600 µl of a migration-inducing media (basal DMEM, 0.1% BSA, 20 ng/µl EGF). Inserts were incubated for 24 hrs at 37°C, 5% CO2, and cells that migrated through the insert into the lower chamber were fixed in methanol, and stained with a solution containing 1X PBS, 0.01 mg/mL DAPI, 1X Triton X 100. Cells that had not migrated through the membrane (top of inserts) were gently scraped off the top of the transwell insert using cotton swabs to ensure quantification of only cell migrated to the bottom of the inserts. The migrated cells were then imaged and counted on an Olympus microscope equipped with the cellSens counting module. All migrated cell counts were normalized to the cell counts from cells containing an empty vector.

### Cell proliferation assay

About 2000 cells per well were seeded in 12-well plates and incubated for 3 hours at 37°C and 5% CO2. Cells were then moved to an Incucyte instrument equipped with a camera and standard cell culture incubator to monitor cell proliferation based on continuous image acquisitions at regular intervals, and proliferation curves were plotted using the Incucyte software.

### Immunohistochemistry

BEAS-2B cells (∼50,000) expressing H2B variant minigenes were seeded in biological triplicates on 35 mm glass bottom plates to ∼50% confluency. Cells were washed 3x with 1X PBS, fixed with 1 mL ice cold methanol for 10 minutes on ice, and then washed 5x with 1X PBS for 5 min. Cells were permeabilized with 1X PBS + 0.2% Triton X100 in 1X PBS for 20 min, and rinsed 3x with 1X PBS. Cells were blocked with a solution containing 1X PBS, 0.01% Triton X100, 10% goat serum for 30 min at 4°C, and then incubated with primary antibodies overnight at 4°C in 1X PBS + 1% BSA. The next day, cells were washed 3x with 1X PBS for 10 min each wash, incubated with secondary antibodies for 1 hr at room temperature, and washed again 3x in 1X PBS for 10 min each wash. Cells were then mounted by adding Prolong-Gold +DAPI dropwise, a coverslip was placed on top of each well, and the coverslip sealed with nail polish. Slides were stored at 4°C in the dark. Images were acquired with a Zeiss 880 confocal microscope and Olympus cellSens Software, and images processed with ImageJ.

### RNA-seq

Total RNA was isolated from ∼1x10^6^ BEAS-2B cells (n=3 biological replicates per group) using the *Quick*-RNA MiniPrep kit (Zymo Research, #R1054; #R1055) as per the manufacture’s protocol. RNA quality and concentration was assessed using an Agilent 4200 Tape Station (Agilent Technologies, # G2991BA) or a QuantiFluor® dsDNA System (Promega, #E2671). Ribosomal RNA was depleted using the QIAseq FastSelect –rRNA HMR Kit (Qiagen, #334376), and the remaining RNA sheared to 300-400 bp before library preparation. Libraries were prepared from 500 ng of total RNA using the KAPA RNA HyperPrep Kit (Kapa Biosystems, #08098107702) and IDT for Illumina TruSeq UD Indexed adapters (Illumina, #20020590). The quality and quantity of the finished libraries were again evaluated using the Agilent 4200 Tape Station (Agilent Technologies, # G2991BA) or QuantiFluor® dsDNA System (Promega, #E2671). Individually indexed libraries were pooled, and 50 bp paired-end sequencing performed on an Illumina NovaSeq 6000 sequencer to an average depth of 84M raw paired-reads per transcriptome. Base calling was done by Illumina RTA3, and the output of NCS was demultiplexed and converted to FastQ format with Illumina Bcl2fastq v1.9.0.

Adaptor sequences and low-quality reads were removed with Trim Galore (v0.6.0) [106]. Trimmed reads were then aligned to the GRCh38 reference genome using STAR (v2.7.8) with parameter “--quantMode GeneCounts” [107]. Differential gene expression (DGE) analyses were conducted using edgeR based on gene counts generated from STAR [108–110], using Benjamini-Hochberg adjusted p-values set to 0.05. ClusterProfiler (v4.0.5) was used to perform gene set enrichment analysis (GSEA) for cancer signatures (C6) and Hallmarks within the Molecular Signatures Database (MSigDB) [111]. Heatmaps of gene expression data were constructed using ComplexHeatmap [112].

### ATAC-seq

Cell lines expressing H2B variant minigenes were grown to 70-90% confluency, and 1x10^5^ cells were cryopreserved (n = 3 biological replicates per group). Cryopreserved cells were sent to Active Motif (Carlsbad, CA) for ATAC-seq assays. Cells were thawed in a 37°C water bath, pelleted, washed with cold PBS, and tagmented as previously described [113], with some modifications based on [114]. In brief, cell pellets were first resuspended in lysis buffer, followed by pelleting and tagmentation using the enzyme and buffer provided in the ATAC-Seq Kit (Active Motif, #53150). Subsequently, tagmented DNA was purified using the MinElute PCR purification kit (Qiagen, #28004), then amplified by PCR for 10 cycles, and further purified using Agencourt AMPure SPRI beads (Beckman Coulter, #B23317). The resulting material was quantified utilizing the iSeq system (Illumina), and subjected to 42-bp paired-end sequencing on a NovaSeq 6000 sequencer (Illumina).

The obtained reads were processed and aligned using the BWA algorithm in “mem mode” with default settings [115]. To ensure data integrity, duplicate reads were removed, and only those reads mapping as matched pairs and uniquely mapped reads with a mapping quality ≥1 were retained for subsequent analysis. Alignments were virtually extended at their 3’-ends to a uniform length of 200 bp, and binned into 32-nt segments across the genome. The resulting histograms (or genomic "signal maps") were stored as bigwig files.

For comparative analysis, we normalized data by down sampling the usable number of tags for each sample within a group to the level of the sample with the fewest usable tags in that group. This procedure ensures an equal number of tags within each group, reflected in the "Used number of tags" column. We used the MACS 2.1.0 algorithm with a p-value cutoff of 1e-7, no control file, and the –nomodel option [116] for peak calling. False peaks (as defined within the ENCODE blacklist [63]) were filtered out, and a FRIP (fraction of reads in peaks) value ≥10% was considered good quality data. Signal maps and peak locations were then used as input data for further analysis with the Active Motif’s proprietary software, which generated detailed Excel tables containing sample comparisons, peak metrics, peak locations, and gene annotations.

Peaks were annotated to the nearest gene using the ‘TxDb.Hsapiens. UCSC.hg38.knownGene’ package v3.16.0, and the ‘annotatePeak’ function in ChIPseeker v1.34.1 with the parameter ‘tssRegion = c(-3000, 500)’ [117]. Separately, peaks were annotated to ENCODE predicted cis-regulatory elements (cCRE) using GenomicRanges v1.50.2 [118] and an overlap of at least 1 bp with ENCODE3 [119] predicted cis-regulatory elements (cCREs). Read coverage at genes and promoters were annotated in GENCODE v33 (hg38) [120], and were plotted using DeepTools v3.5.2 [121]. Sample-specific bigwig files were averaged across replicates using the ‘bigwigAverage’ function with the parameters, ‘--binSize 10 -of “bigwig”’. For gene body plots, the ‘computeMatrix’ function was run with the parameters ‘scale-regions -- regionBodyLength 5000 -a 3000 -b 3000 --missingDataAsZero --binSize 10 --transcriptID gene - -transcript_id_designator gene_id’, and specifying the ENCODE Unified GRCh38 Blacklist (https://www.encodeproject.org/files/ENCFF356LFX/) using the ‘-bl’ parameter. The function ‘plotHeatmap’ was run to generate an associated image. For the heatmaps centered on the transcription start site (TSS), the workflow was the same except the ‘computeMatrix’ parameters ‘scale-regions --regionBodyLength 5000’ were replaced with ‘reference-point --referencePoint TSS’.

Gene ontology and pathway enrichment analysis were performed on differentially accessible peaks (*p_adj_* ≤ 0.1), and were separated into up-or down-regulated lists. Only peaks overlapping promoters were retained and grouped together if they were annotated to the same nearest gene. The resulting gene lists were used for over-representation analysis using the ‘compareCluster’ function in clusterProfiler v4.6.2 with a default cutoff of *p_adj_* ≤ 0.05 [94]. MSigDB annotations were retrieved using msigdbr v7.5.1 [122, 123], and summary plots of the enrichment results were created with enrichplot v1.18.4. Finally, genes with differentially accessible promoters (determined as described above for the pathway analyses) were compared with the lists of RNA-seq differentially expressed genes (*p_adj_* cutoff ≤ 0.05). These comparisons were plotted as Venn diagrams using the ggVennDiagram v1.2.3 package [124].

## Acknowledgements

We are grateful to the Fondufe-Mittendorf lab for scientific discussions contributing to this manuscript and members of the VAI Core Facilities (Genomics, Optical Imaging, Flow Cytometry as well as Bioinformatics & Biostatistics) for technical and data analyses assistance. RLB is supported by R50CA293837. JDL is supported by R01CA266078. YFM is supported by the National Science Foundation grant NSF/MCB 016515, National Institutes of Health grants R01ES024478, R01ES034253, and 1R01ES036051-01, and the Van Andel Institute (VAI).

## Author Contributions

Conceptualization: WNS and YFM; investigation: WNS, YAH, KC, ME, RNC, HD, RLB, DPM, MGP, YD, JDL and YFM; data analysis: WNS, YAH, KC, DPM, RNC, HD, EL, ZF, KL and YFM; data curation: WNS, YH, KC, ME, RNC, HD and RLB; writing—original draft preparation: WNS and YFM; writing-editing: WNS, DPC, RLB, DPM, MGP, YD, JDL and YFM; visualization: WNS, YH, DPM, DPC, and YFM; resources: MGP, YD, JDL and YFM; supervision: YFM; funding acquisition: YFM. All authors have read and agreed to the published version of the manuscript.

## References

1. Hsu, C.J., et al., The Role of MacroH2A Histone Variants in Cancer. Cancers (Basel), 2021. 13(12).

2. Mariño-Ramírez, L., et al., The Histone Database: an integrated resource for histones and histone fold-containing proteins. Database (Oxford), 2011. 2011: p. bar048.

3. Luger, K., et al., Crystal structure of the nucleosome core particle at 2.8 A resolution. Nature, 1997. 389(6648): p. 251-60.

4. Hergeth, S.P. and R. Schneider, The H1 linker histones: multifunctional proteins beyond the nucleosomal core particle. EMBO Rep, 2015. 16(11): p. 1439–53.

5. Bonenfant, D., et al., Characterization of histone H2A and H2B variants and their post-translational modifications by mass spectrometry. Mol Cell Proteomics, 2006. 5(3): p. 541–52.

6. Flavahan, W.A., E. Gaskell, and B.E. Bernstein, Epigenetic plasticity and the hallmarks of cancer. Science, 2017. 357(6348).

7. Schwartzentruber, J., et al., Driver mutations in histone H3.3 and chromatin remodelling genes in paediatric glioblastoma. Nature, 2012. 482(7384): p. 226-31.

8. Taylor, K.R., et al., Recurrent activating ACVR1 mutations in diffuse intrinsic pontine glioma. Nat Genet, 2014. 46(5): p. 457–461.

9. Bagert, J.D., et al., Oncohistone mutations enhance chromatin remodeling and alter cell fates. Nat Chem Biol, 2021. 17(4): p. 403–411.

10. Nacev, B.A., et al., The expanding landscape of ’oncohistone’ mutations in human cancers. Nature, 2019. 567(7749): p. 473-478.

11. Schaefer, I.M., et al., Immunohistochemistry for histone H3G34W and H3K36M is highly specific for giant cell tumor of bone and chondroblastoma, respectively, in FNA and core needle biopsy. Cancer Cytopathol, 2018. 126(8): p. 552–566.

12. Behjati, S., et al., Distinct H3F3A and H3F3B driver mutations define chondroblastoma and giant cell tumor of bone. Nat Genet, 2013. 45(12): p. 1479–82.

13. Papillon-Cavanagh, S., et al., Impaired H3K36 methylation defines a subset of head and neck squamous cell carcinomas. Nat Genet, 2017. 49(2): p. 180–185.

14. Yusufova, N., et al., Histone H1 loss drives lymphoma by disrupting 3D chromatin architecture. Nature, 2021. 589(7841): p. 299-305.

15. Bennett, R.L., et al., A Mutation in Histone H2B Represents a New Class of Oncogenic Driver. Cancer Discov, 2019.

16. Maze, I., et al., Every amino acid matters: essential contributions of histone variants to mammalian development and disease. Nat Rev Genet, 2014. 15(4): p. 259–71.

17. Skene, P.J. and S. Henikoff, Histone variants in pluripotency and disease. Development, 2013. 140(12): p. 2513–24.

18. Wang, T., et al., Histone variants: critical determinants in tumour heterogeneity. Front Med, 2019. 13(3): p. 289–297.

19. Chew, G.L., et al., Short H2A histone variants are expressed in cancer. Nat Commun, 2021. 12(1): p. 490.

20. Talbert, P.B. and S. Henikoff, Histone variants at a glance. J Cell Sci, 2021. 134(6).

21. Espinoza Pereira, K.N., et al., Histone mutations in cancer. Biochem Soc Trans, 2023. 51(5): p. 1749–1763.

22. Weber, C.M. and S. Henikoff, Histone variants: dynamic punctuation in transcription. Genes Dev, 2014. 28(7): p. 672–82.

23. Talbert, P.B. and S. Henikoff, Histone variants--ancient wrap artists of the epigenome. Nat Rev Mol Cell Biol, 2010. 11(4): p. 264–75.

24. Zlatanova, J., et al., The nucleosome family: dynamic and growing. Structure, 2009. 17(2): p. 160–71.

25. Saintilnord, W.N. and Y. Fondufe-Mittendorf, Arsenic-induced epigenetic changes in cancer development. Semin Cancer Biol, 2021. 76: p. 195–205.

26. Corujo, D. and M. Buschbeck, Post-Translational Modifications of H2A Histone Variants and Their Role in Cancer. Cancers (Basel), 2018. 10(3).

27. Talbert, P.B. and S. Henikoff, Histone variants on the move: substrates for chromatin dynamics. Nat Rev Mol Cell Biol, 2017. 18(2): p. 115–126.

28. Vardabasso, C., et al., Histone variants: emerging players in cancer biology. Cell Mol Life Sci, 2014. 71(3): p. 379–404.

29. Zink, L.M. and S.B. Hake, Histone variants: nuclear function and disease. Curr Opin Genet Dev, 2016. 37: p. 82–89.

30. Vieira-Silva, T.S., et al., Histone variant MacroH2A1 is downregulated in prostate cancer and influences malignant cell phenotype. Cancer Cell Int, 2019. 19: p. 112.

31. Kallappagoudar, S., et al., Histone H3 mutations--a special role for H3.3 in tumorigenesis? Chromosoma, 2015. 124(2): p. 177–89.

32. Weyemi, U., et al., The histone variant H2A.X is a regulator of the epithelial– mesenchymal transition. Nature Communications, 2016. 7(1): p. 10711.

33. Svotelis, A., et al., H2A.Z overexpression promotes cellular proliferation of breast cancer cells. Cell Cycle, 2010. 9(2): p. 364–70.

34. Yang, H.D., et al., Oncogenic potential of histone-variant H2A.Z.1 and its regulatory role in cell cycle and epithelial-mesenchymal transition in liver cancer. Oncotarget, 2016. 7(10): p. 11412-23.

35. Sun, Z. and E. Bernstein, Histone variant macroH2A: from chromatin deposition to molecular function. Essays Biochem, 2019. 63(1): p. 59–74.

36. Dhahri, H., et al., Beyond the Usual Suspects: Examining the Role of Understudied Histone Variants in Breast Cancer. Int J Mol Sci, 2024. 25(12).

37. Ghiraldini, F.G., D. Filipescu, and E. Bernstein, Solid tumours hijack the histone variant network. Nat Rev Cancer, 2021. 21(4): p. 257–275.

38. Weinstein, J.N., et al., The Cancer Genome Atlas Pan-Cancer analysis project. Nat Genet, 2013. 45(10): p. 1113–20.

39. Mohammad, F. and K. Helin, Oncohistones: drivers of pediatric cancers. Genes Dev, 2017. 31(23-24): p. 2313–2324.

40. Sturm, D., et al., Hotspot mutations in H3F3A and IDH1 define distinct epigenetic and biological subgroups of glioblastoma. Cancer Cell, 2012. 22(4): p. 425–37.

41. Toledo, R.A., et al., Recurrent Mutations of Chromatin-Remodeling Genes and Kinase Receptors in Pheochromocytomas and Paragangliomas. Clin Cancer Res, 2016. 22(9): p. 2301–10.

42. Wan, Y.C.E., et al., Cancer-associated histone mutation H2BG53D disrupts DNA-histone octamer interaction and promotes oncogenic phenotypes. Signal Transduct Target Ther, 2020. 5(1): p. 27.

43. Wan, Y.C.E., et al., The H2BG53D oncohistone directly upregulates ANXA3 transcription and enhances cell migration in pancreatic ductal adenocarcinoma. Signal Transduct Target Ther, 2020. 5(1): p. 106.

44. Kalashnikova, A.A., et al., The role of the nucleosome acidic patch in modulating higher order chromatin structure. J R Soc Interface, 2013. 10(82): p. 20121022.

45. Valencia, A.M., et al., Recurrent SMARCB1 Mutations Reveal a Nucleosome Acidic Patch Interaction Site That Potentiates mSWI/SNF Complex Chromatin Remodeling. Cell, 2019. 179(6): p. 1342–1356.e23.

46. Wan, Y.C.E. and K.M. Chan, Histone H2B Mutations in Cancer. Biomedicines, 2021. 9(6).

47. Worden, E.J., et al., Mechanism of Cross-talk between H2B Ubiquitination and H3 Methylation by Dot1L. Cell, 2019. 176(6): p. 1490–1501.e12.

48. Tsunaka, Y., et al., Alteration of the nucleosomal DNA path in the crystal structure of a human nucleosome core particle. Nucleic Acids Res, 2005. 33(10): p. 3424–34.

49. Polach, K.J. and J. Widom, Mechanism of protein access to specific DNA sequences in chromatin: a dynamic equilibrium model for gene regulation. J Mol Biol, 1995. 254(2): p. 130–49.

50. Li, G. and J. Widom, Nucleosomes facilitate their own invasion. Nat Struct Mol Biol, 2004. 11(8): p. 763–9.

51. Polach, K.J. and J. Widom, A model for the cooperative binding of eukaryotic regulatory proteins to nucleosomal target sites. J Mol Biol, 1996. 258(5): p. 800–12.

52. Miller, J.A. and J. Widom, Collaborative competition mechanism for gene activation in vivo. Mol Cell Biol, 2003. 23(5): p. 1623–32.

53. Rea, M., et al., Quantitative Mass Spectrometry Reveals Changes in Histone H2B Variants as Cells Undergo Inorganic Arsenic-Mediated Cellular Transformation. Mol Cell Proteomics, 2016. 15(7): p. 2411–22.

54. Luo, Y., et al., Nucleosomes accelerate transcription factor dissociation. Nucleic Acids Res, 2014. 42(5): p. 3017–27.

55. Luo, Y., J.A. North, and M.G. Poirier, Single molecule fluorescence methodologies for investigating transcription factor binding kinetics to nucleosomes and DNA. Methods, 2014. 70(2-3): p. 108–18.

56. Zlatanova, J., C. Seebart, and M. Tomschik, The linker-protein network: control of nucleosomal DNA accessibility. Trends Biochem Sci, 2008. 33(6): p. 247–53.

57. Ngo, T.T., et al., Asymmetric unwrapping of nucleosomes under tension directed by DNA local flexibility. Cell, 2015. 160(6): p. 1135–44.

58. Mondal, A. and A.B. Kolomeisky, Why Are Nucleosome Breathing Dynamics Asymmetric? J Phys Chem Lett, 2024. 15(2): p. 422–431.

59. García, A., et al., Asymmetrical nucleosomal DNA signatures regulate transcriptional directionality. Cell Rep, 2023. 43(1): p. 113605.

60. Simon, M., et al., Histone fold modifications control nucleosome unwrapping and disassembly. Proc Natl Acad Sci U S A, 2011. 108(31): p. 12711–6.

61. Walkiewicz, M.P., et al., Tracking histone variant nucleosomes across the human cell cycle using biophysical, biochemical, and cytological analyses. Methods Mol Biol, 2014. 1170: p. 589–615.

62. Feierman, E.R., et al., Histone variant H2BE enhances chromatin accessibility in neurons to promote synaptic gene expression and long-term memory. Mol Cell, 2024. 84(15): p. 2822–2837.e11.

63. An integrated encyclopedia of DNA elements in the human genome. Nature, 2012. 489(7414): p. 57-74.

64. Reyes, M., et al., Simultaneous profiling of gene expression and chromatin accessibility in single cells. Adv Biosyst, 2019. 3(11).

65. Starks, R.R., et al., Combined analysis of dissimilar promoter accessibility and gene expression profiles identifies tissue-specific genes and actively repressed networks. Epigenetics Chromatin, 2019. 12(1): p. 16.

66. Landschulz, W.H., P.F. Johnson, and S.L. McKnight, The leucine zipper: a hypothetical structure common to a new class of DNA binding proteins. Science, 1988. 240(4860): p. 1759-64.

67. Kerppola, T.K. and T. Curran, Fos-Jun heterodimers and Jun homodimers bend DNA in opposite orientations: implications for transcription factor cooperativity. Cell, 1991. 66(2): p. 317–26.

68. Shaulian, E. and M. Karin, AP-1 as a regulator of cell life and death. Nat Cell Biol, 2002. 4(5): p. E131–6.

69. Eferl, R. and E.F. Wagner, AP-1: a double-edged sword in tumorigenesis. Nat Rev Cancer, 2003. 3(11): p. 859–68.

70. Wei, D., et al., Drastic down-regulation of Krüppel-like factor 4 expression is critical in human gastric cancer development and progression. Cancer Res, 2005. 65(7): p. 2746–54.

71. Yao, J., et al., miR-450b-3p inhibited the proliferation of gastric cancer via regulating KLF7. Cancer Cell Int, 2020. 20: p. 47.

72. Zou, H., et al., ATXN3 promotes breast cancer metastasis by deubiquitinating KLF4. Cancer Lett, 2019. 467: p. 19–28.

73. Gupta, R., et al., KLF7 promotes pancreatic cancer growth and metastasis by up-regulating ISG expression and maintaining Golgi complex integrity. Proc Natl Acad Sci U S A, 2020. 117(22): p. 12341–12351.

74. Kim, S., N.K. Yu, and B.K. Kaang, CTCF as a multifunctional protein in genome regulation and gene expression. Exp Mol Med, 2015. 47: p. e166.

75. Phillips, J.E. and V.G. Corces, CTCF: master weaver of the genome. Cell, 2009. 137(7): p. 1194–211.

76. Hua, S., et al., Genomic analysis of estrogen cascade reveals histone variant H2A.Z associated with breast cancer progression. Mol Syst Biol, 2008. 4: p. 188.

77. Sporn, J.C. and B. Jung, Differential regulation and predictive potential of MacroH2A1 isoforms in colon cancer. Am J Pathol, 2012. 180(6): p. 2516–26.

78. Sporn, J.C., et al., Histone macroH2A isoforms predict the risk of lung cancer recurrence. Oncogene, 2009. 28(38): p. 3423–8.

79. Ragusa, D. and P. Vagnarelli, Contribution of histone variants to aneuploidy: a cancer perspective. Front Genet, 2023. 14: p. 1290903.

80. Tóth, K., et al., Histone-and DNA sequence-dependent stability of nucleosomes studied by single-pair FRET. Cytometry A, 2013. 83(9): p. 839–46.

81. Chua, E.Y., et al., The mechanics behind DNA sequence-dependent properties of the nucleosome. Nucleic Acids Res, 2012. 40(13): p. 6338–52.

82. Bilokapic, S., M. Strauss, and M. Halic, Histone octamer rearranges to adapt to DNA unwrapping. Nat Struct Mol Biol, 2018. 25(1): p. 101–108.

83. Ávila-López, P.A., et al., H2A.Z overexpression suppresses senescence and chemosensitivity in pancreatic ductal adenocarcinoma. Oncogene, 2021. 40(11): p. 2065–2080.

84. Damdindorj, L., et al., A comparative analysis of constitutive promoters located in adeno-associated viral vectors. PLoS One, 2014. 9(8): p. e106472.

85. Singh, R.K., et al., Excess histone levels mediate cytotoxicity via multiple mechanisms. Cell Cycle, 2010. 9(20): p. 4236–44.

86. Choi, C.R., et al., Clonal evolution of colorectal cancer in IBD. Nat Rev Gastroenterol Hepatol, 2017. 14(4): p. 218–229.

87. Takeshima, H. and T. Ushijima, Accumulation of genetic and epigenetic alterations in normal cells and cancer risk. NPJ Precis Oncol, 2019. 3: p. 7.

88. Lu, R., et al., Epigenetic Perturbations by Arg882-Mutated DNMT3A Potentiate Aberrant Stem Cell Gene-Expression Program and Acute Leukemia Development. Cancer Cell, 2016. 30(1): p. 92–107.

89. Langille, E., et al., Loss of Epigenetic Regulation Disrupts Lineage Integrity, Induces Aberrant Alveogenesis, and Promotes Breast Cancer. Cancer Discov, 2022. 12(12): p. 2930–2953.

90. Vicente-Dueñas, C., et al., Epigenetic Priming in Cancer Initiation. Trends Cancer, 2018. 4(6): p. 408–417.

91. Vaz, M., et al., Chronic Cigarette Smoke-Induced Epigenomic Changes Precede Sensitization of Bronchial Epithelial Cells to Single-Step Transformation by KRAS Mutations. Cancer Cell, 2017. 32(3): p. 360–376.e6.

92. Wilks, C., et al., recount3: summaries and queries for large-scale RNA-seq expression and splicing. Genome Biol, 2021. 22(1): p. 323.

93. Love, M.I., W. Huber, and S. Anders, Moderated estimation of fold change and dispersion for RNA-seq data with DESeq2. Genome Biol, 2014. 15(12): p. 550.

94. Yu, G., et al., clusterProfiler: an R package for comparing biological themes among gene clusters. Omics, 2012. 16(5): p. 284–7.

95. Therneau, T.M. A Package for Survival Analysis in R. 2023.

96. Therneau, P.M.G.a.T.M., Modeling Survival Data: Extending the Cox Model. 2000: Springer.

97. Liang, S.D., et al., DNA sequence preferences of GAL4 and PPR1: how a subset of Zn2 Cys6 binuclear cluster proteins recognizes DNA. Mol Cell Biol, 1996. 16(7): p. 3773–80.

98. Anderson, J.D. and J. Widom, Sequence and position-dependence of the equilibrium accessibility of nucleosomal DNA target sites. J Mol Biol, 2000. 296(4): p. 979–87.

99. Bernier, M., et al., Linker histone H1 and H3K56 acetylation are antagonistic regulators of nucleosome dynamics. Nat Commun, 2015. 6: p. 10152.

100. Burge, N.L., et al., H1.0 C Terminal Domain Is Integral for Altering Transcription Factor Binding within Nucleosomes. Biochemistry, 2022. 61(8): p. 625–638.

101. Luger, K., T.J. Rechsteiner, and T.J. Richmond, Preparation of nucleosome core particle from recombinant histones. Methods Enzymol, 1999. 304: p. 3–19.

102. Shimko, J.C., et al., Preparation of fully synthetic histone H3 reveals that acetyl-lysine 56 facilitates protein binding within nucleosomes. J Mol Biol, 2011. 408(2): p. 187–204.

103. Schwarz, P.M., et al., Reversible oligonucleosome self-association: dependence on divalent cations and core histone tail domains. Biochemistry, 1996. 35(13): p. 4009–15.

104. Clegg, R.M., Fluorescence resonance energy transfer and nucleic acids. Methods Enzymol, 1992. 211: p. 353–88.

105. Yang, N., et al., A hyper-quiescent chromatin state formed during aging is reversed by regeneration. Mol Cell, 2023. 83(10): p. 1659–1676.e11.

106. Martin, M., Cutadapt Removes Adapter Sequences From High-Throughput Sequencing Reads. EMBnet.journal, 2011. 17(1).

107. Dobin, A., et al., STAR: ultrafast universal RNA-seq aligner. Bioinformatics, 2013. 29(1): p. 15–21.

108. Robinson, M.D., D.J. McCarthy, and G.K. Smyth, edgeR: a Bioconductor package for differential expression analysis of digital gene expression data. Bioinformatics, 2010. 26(1): p. 139–40.

109. McCarthy, D.J., Y. Chen, and G.K. Smyth, Differential expression analysis of multifactor RNA-Seq experiments with respect to biological variation. Nucleic Acids Res, 2012. 40(10): p. 4288–97.

110. Chen, Y., A.T. Lun, and G.K. Smyth, From reads to genes to pathways: differential expression analysis of RNA-Seq experiments using Rsubread and the edgeR quasi-likelihood pipeline. F1000Res, 2016. 5: p. 1438.

111. Wu, T., et al., clusterProfiler 4.0: A universal enrichment tool for interpreting omics data. Innovation (Camb), 2021. 2(3): p. 100141.

112. Gu, Z., R. Eils, and M. Schlesner, Complex heatmaps reveal patterns and correlations in multidimensional genomic data. Bioinformatics, 2016. 32(18): p. 2847–9.

113. Schep, A.N., et al., Structured nucleosome fingerprints enable high-resolution mapping of chromatin architecture within regulatory regions. Genome Res, 2015. 25(11): p. 1757–70.

114. Corces, M.R., et al., An improved ATAC-seq protocol reduces background and enables interrogation of frozen tissues. Nat Methods, 2017. 14(10): p. 959–962.

115. Li, H. and R. Durbin, Fast and accurate long-read alignment with Burrows-Wheeler transform. Bioinformatics, 2010. 26(5): p. 589–95.

116. Zhang, Y., et al., Model-based analysis of ChIP-Seq (MACS). Genome Biol, 2008. 9(9): p. R137.

117. Yu, G., L.G. Wang, and Q.Y. He, ChIPseeker: an R/Bioconductor package for ChIP peak annotation, comparison and visualization. Bioinformatics, 2015. 31(14): p. 2382–3.

118. Lawrence, M., et al., Software for computing and annotating genomic ranges. PLoS Comput Biol, 2013. 9(8): p. e1003118.

119. Moore, J.E., et al., Expanded encyclopaedias of DNA elements in the human and mouse genomes. Nature, 2020. 583(7818): p. 699-710.

120. Frankish, A., et al., GENCODE 2021. Nucleic Acids Res, 2021. 49(D1): p. D916-d923.

121. Ramírez, F., et al., deepTools2: a next generation web server for deep-sequencing data analysis. Nucleic Acids Res, 2016. 44(W1): p. W160–5.

122. Subramanian, A., et al., Gene set enrichment analysis: a knowledge-based approach for interpreting genome-wide expression profiles. Proc Natl Acad Sci U S A, 2005. 102(43): p. 15545–50.

123. Liberzon, A., et al., The Molecular Signatures Database (MSigDB) hallmark gene set collection. Cell Syst, 2015. 1(6): p. 417–425.

124. Gao, C.H., G. Yu, and P. Cai, ggVennDiagram: An Intuitive, Easy-to-Use, and Highly Customizable R Package to Generate Venn Diagram. Front Genet, 2021. 12: p. 706907.

