## Supplemental Figures for "Aberrant expression of histone H2B variants reshape chromatin and alter oncogenic gene expression programs"

**Supplemental Figures**
Saintilnord et al. 2024

**Suppl Fig 1**. Histone H2B genes are differentially expressed across cancers.

**Suppl Fig 2**. Known oncogenes and chromatin remodelers are co-expressed with H2B variants. Images are broken across multiple pages.

**Suppl Fig 3**. Differentially co-expressed C6 oncogenic signatures for patients who over-expressed an H2B variant that is associated with poor patient survival.

**Suppl Fig 4**. Differentially co-expressed Hallmarks of H2B variants significantly associated with survival.

**Suppl Fig 5**. H2B variants exhibit unique mutational hotspots.

**Suppl Fig 6**. H2B variants alter nucleosome structure and DNA accessibility.

**Suppl Fig 7**. Histone H2B variants alter access to nucleosome linker DNA.

**Suppl Fig 8**. H2B variant nucleosomes form organized nucleosome arrays.

**Suppl Fig 9**. Histone H2B variants are expressed and incorporated into chromatin.

**Suppl Fig 10**. Histone H2B variants alter chromatin accessibility.

**Suppl Fig 11**. Histone H2B variants alter gene expression programs. Images are broken across multiple pages.

**Suppl Fig 12**. H2B variants (by themselves) do not significantly alter cell migration or proliferation.

**Suppl Fig 13**. H2B variant DEGs and their association with DARs.

[Remainder of page intentionally left blank]

**Suppl Fig 1: Histone H2B genes are differentially expressed across cancers.** A) Heatmap of the expression of H2B genes across TCGA cancer types relative to their normal adjacent tissues. **B**) Same as in **A**, but H2B gene expression was normalized to *H2BC4* expression. **C**) Survival curves for those patients overexpressing H2B genes that are significantly associated with poor prognosis using the Cox proportional hazard regression model. Shaded area around each line represents the 95% confidence interval. As expected, older patients (in the 75$^{th}$ percentile) had lower overall survival rates.

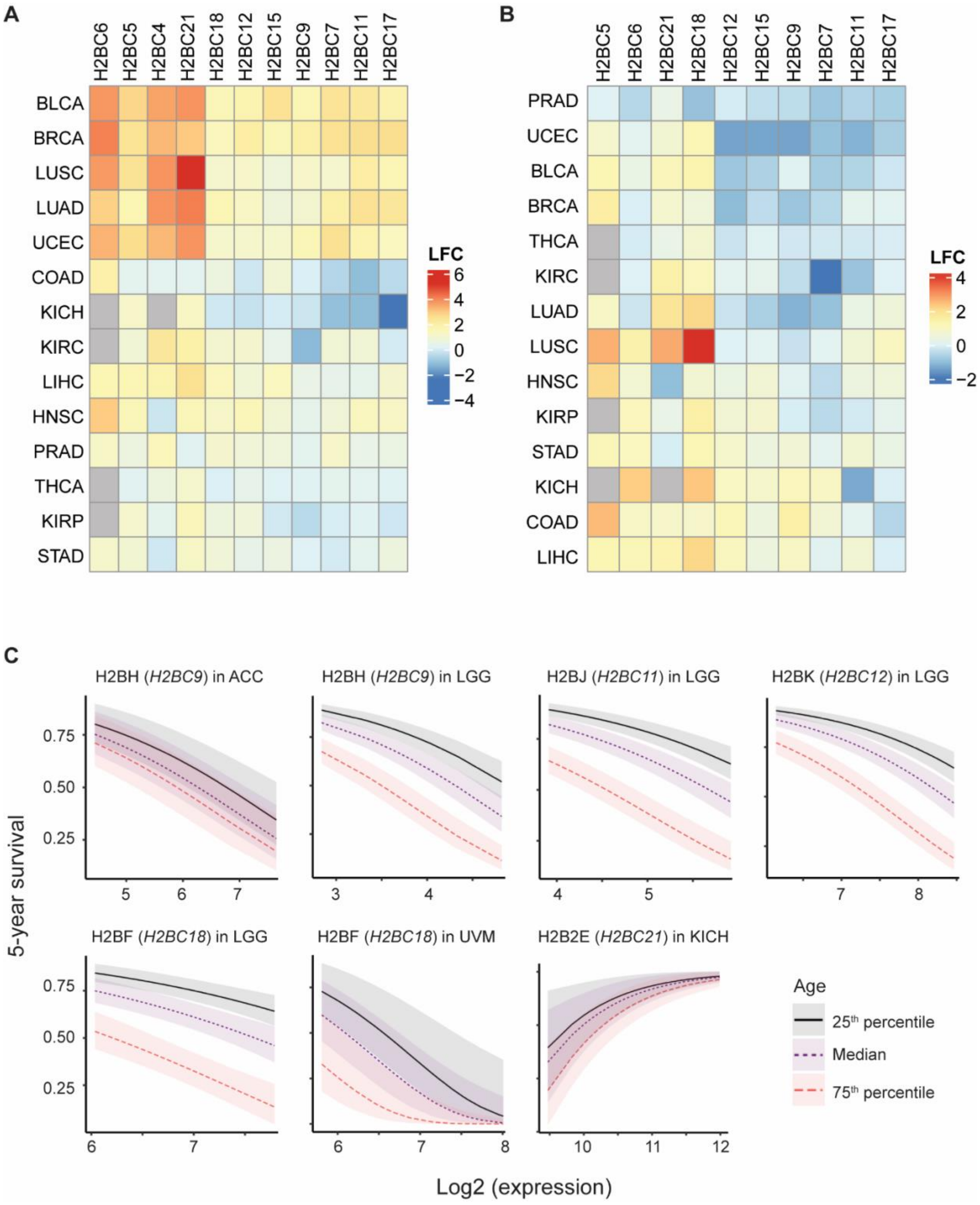

Suppl Fig 2

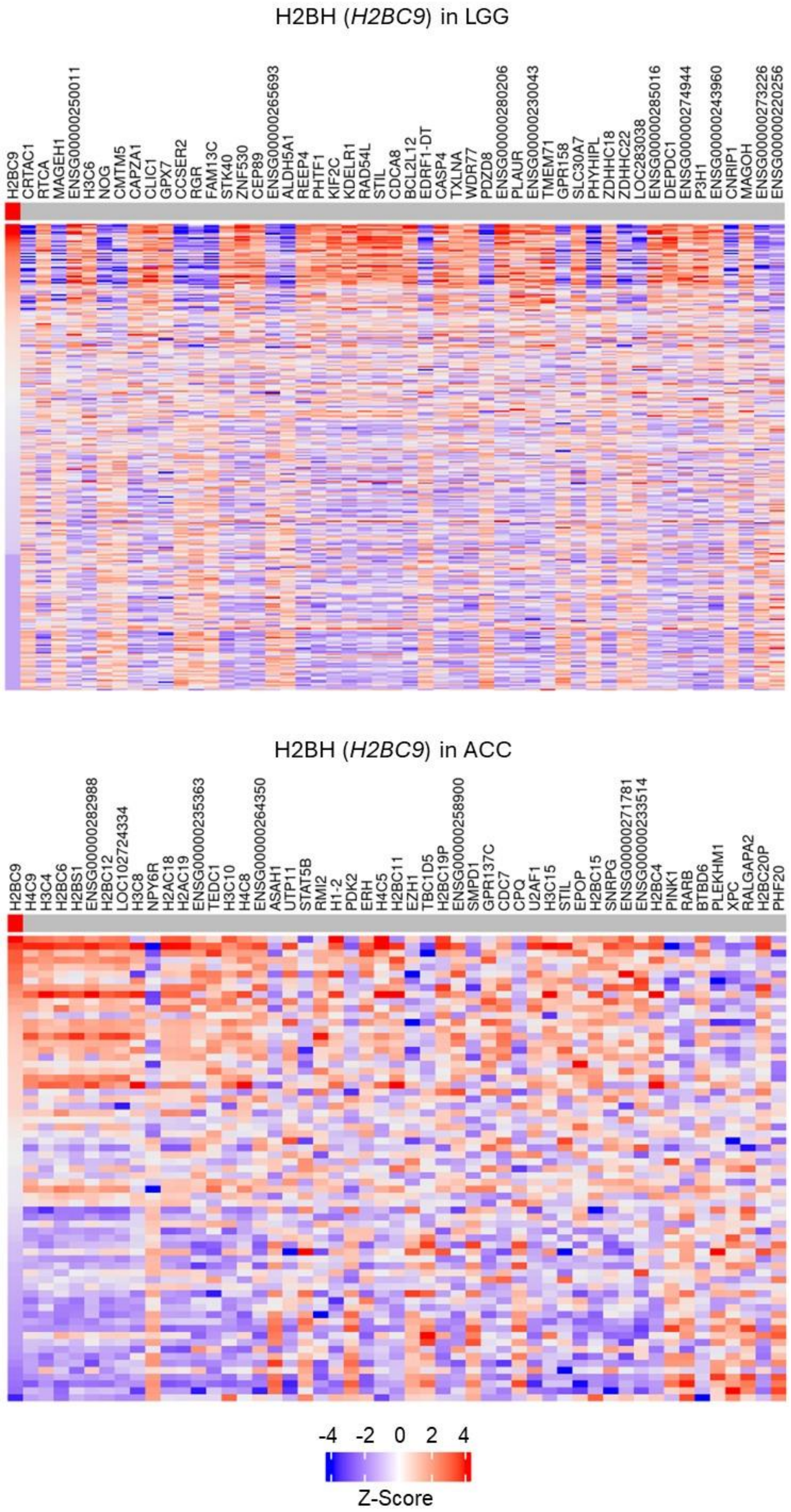

Suppl Fig 2, cont.

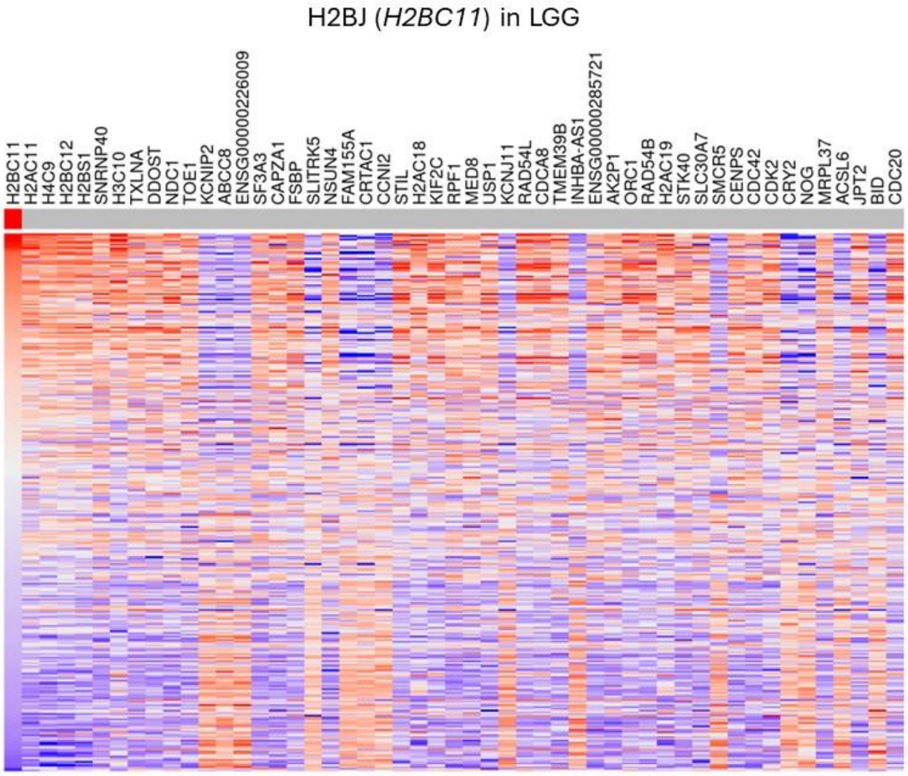


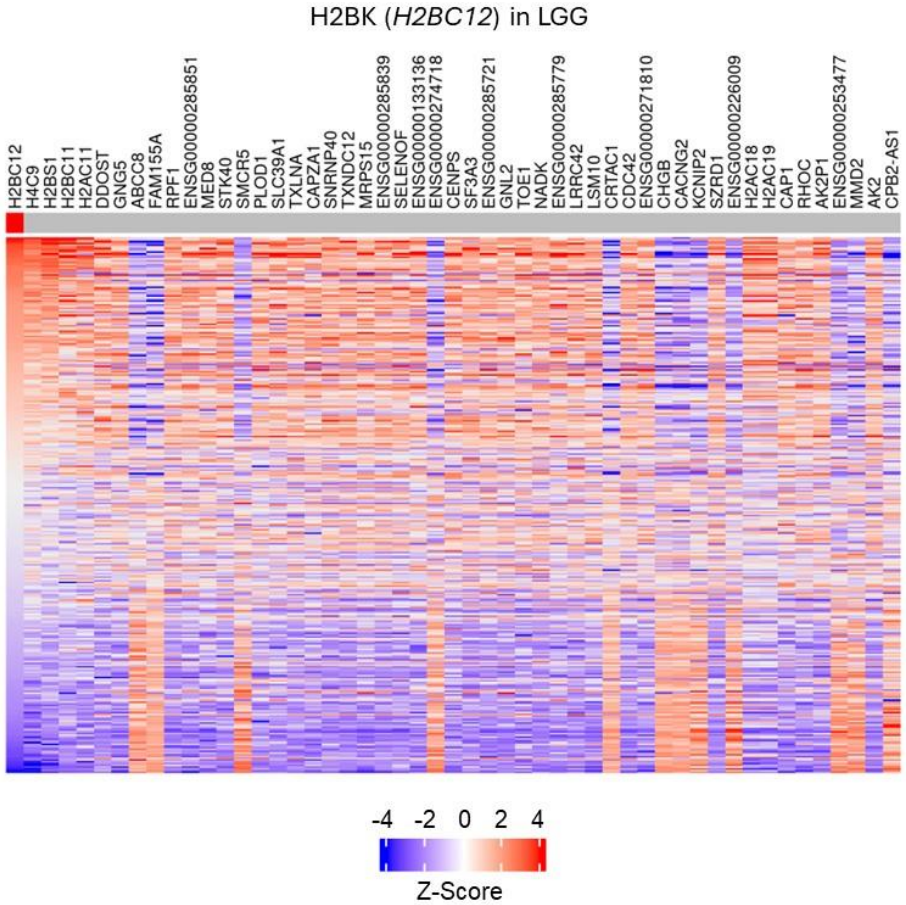

Suppl Fig 2, cont.

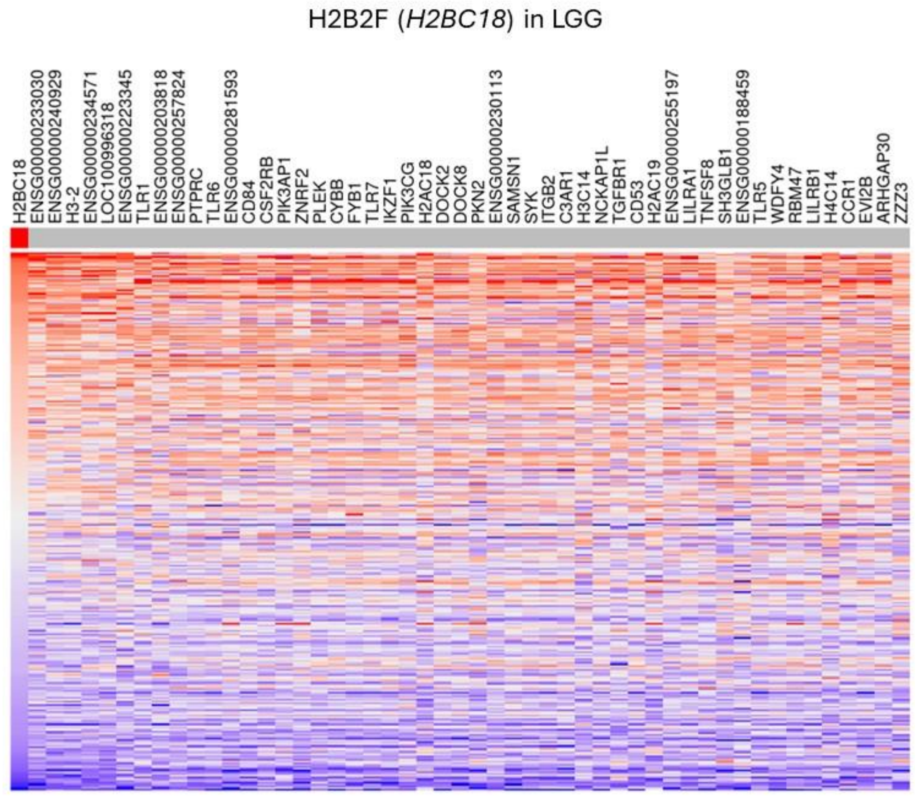


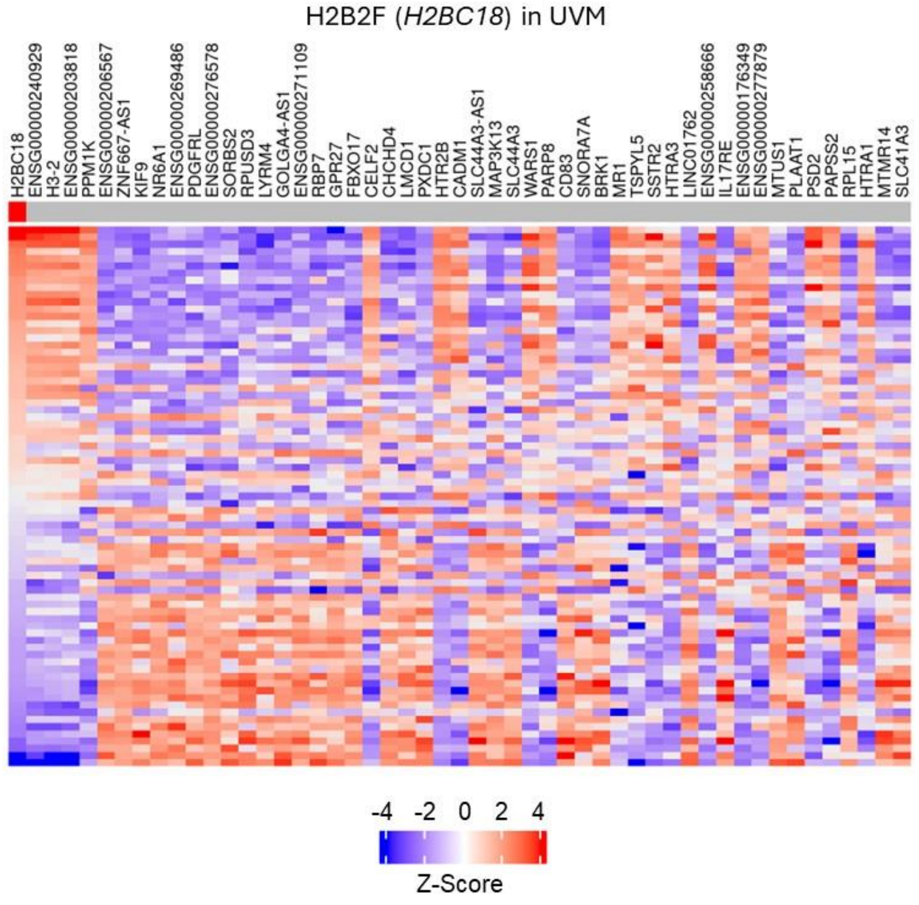

Suppl Fig 2, cont.

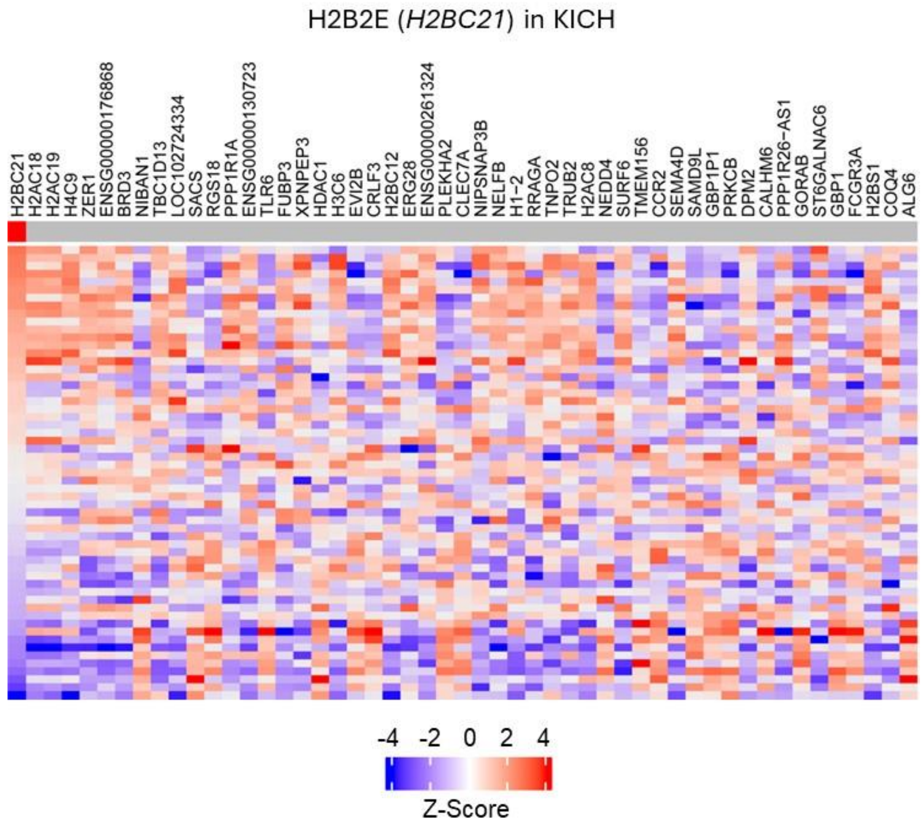


**Suppl Fig 2. Known oncogenes and chromatin remodelers are coexpress with H2B variants.** Shown are the top 50 differentially co-expressed genes for patients who over-expressed an H2B variant that is associated with poor patient survival. Each column is a single patient. LGG = low grade glioma; ACC = adenoid cystic carcinoma; UVM = uveal melanoma; KICH=chromophobe renal cell carcinoma.

[Remainder of page intentionally left blank]

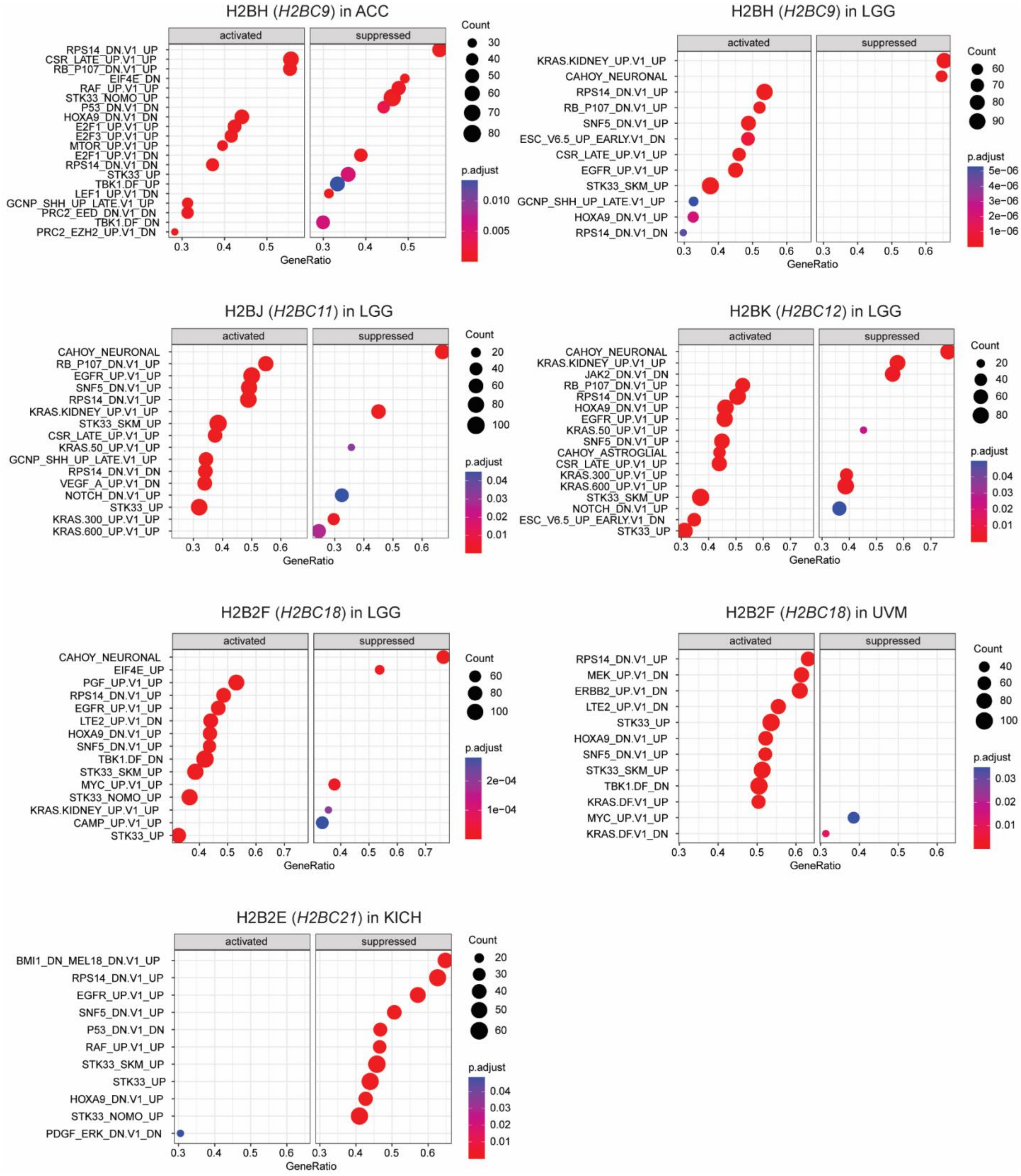


**Suppl Fig 3. Differentially co-expressed C6 oncogenic signatures for patients who overexpress an H2B variant that is associated with poor patient survival.** Gene names are in parentheses. LGG = low grade glioma; ACC = adenoid cystic carcinoma; UVM = uveal melanoma; KICH=chromophobe renal cell carcinoma.

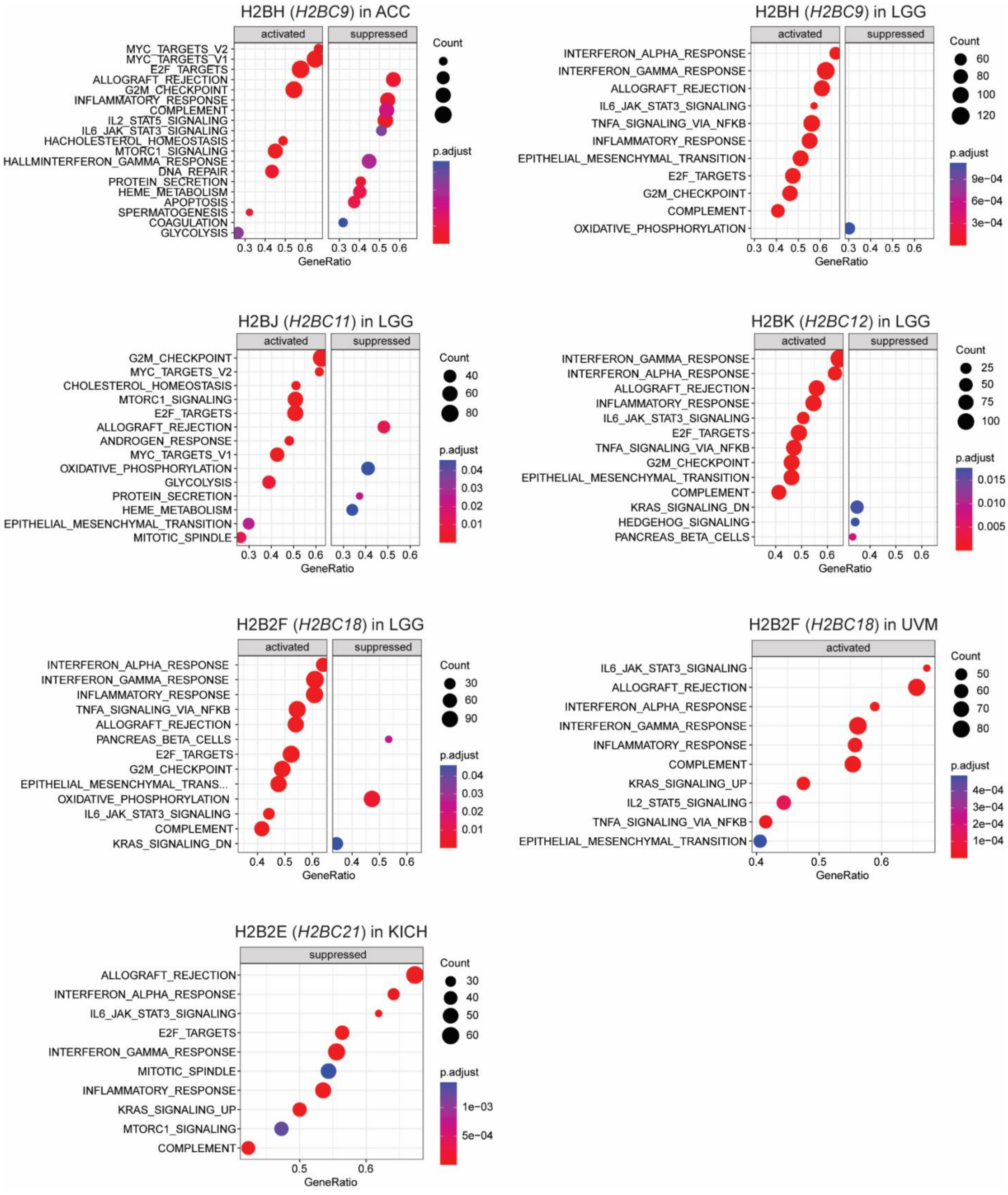


**Suppl Fig 4. Differentially co-expressed Hallmarks of H2B variants significantly associated with survival.** Gene names are in parentheses. LGG = low grade glioma; ACC = adenoid cystic carcinoma; UVM = uveal melanoma; KICH=chromophobe renal cell carcinoma.

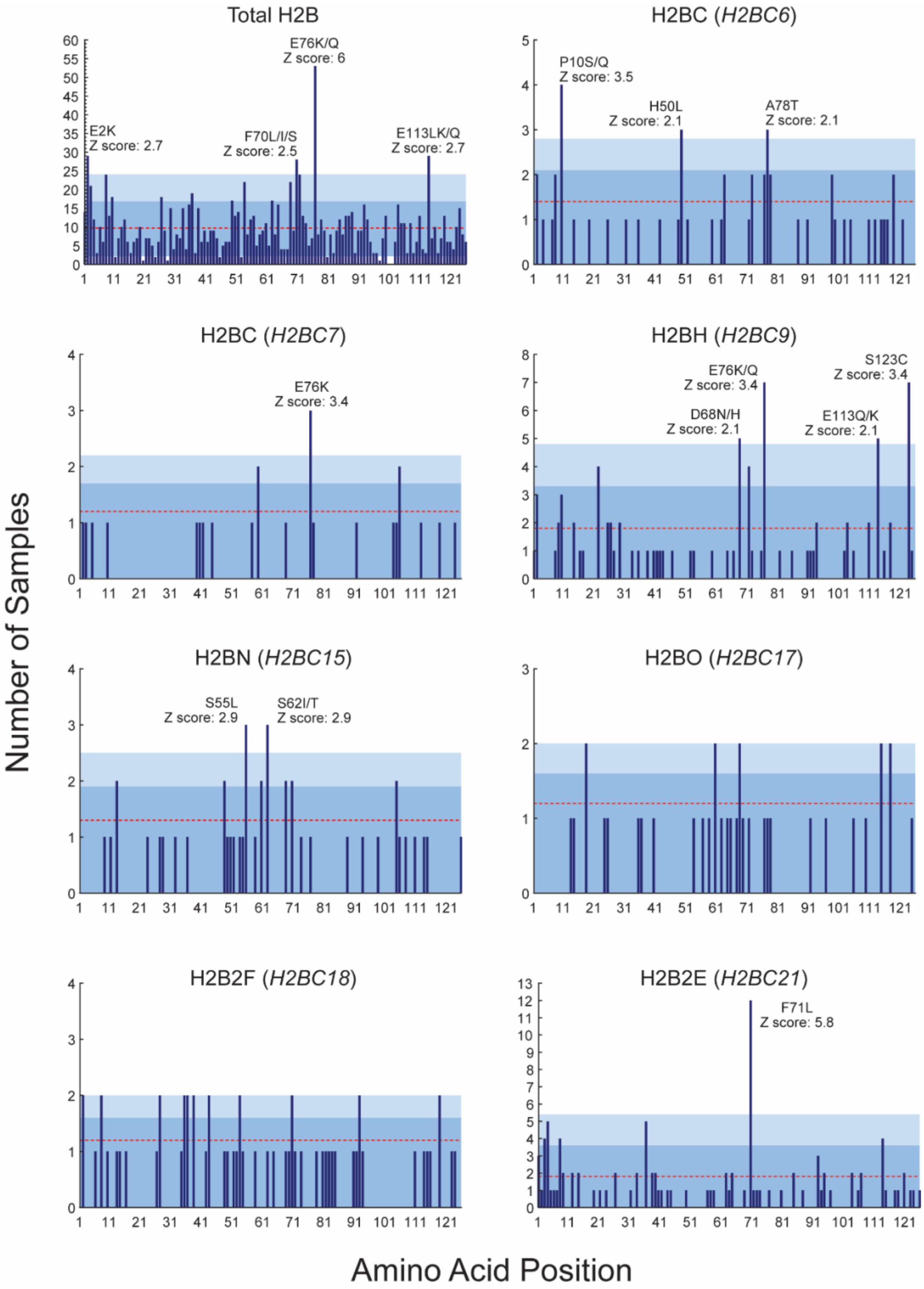


**Suppl Fig 5. H2B variants exhibit unique mutational hotspots**. H2B missense mutations were derived from 65,489 non-redundant patient samples in the cBioPortal for Cancer Genomics database, and encompass all cancers. Frequently occurring mutations are those with a z-score >2, and are labeled in each panel. The gene encoding each H2B variant is indicated in parentheses. Red lines are the average number of mutations for each histone variant, with dark and light blue shading representing the first and second standard deviations from the average, respectively. "Total H2B" is a cross-cancer mutation summary for all H2B variants combined, and shows the same H2BE76K/Q hotspot previously identified by Bennett et al. 2019.

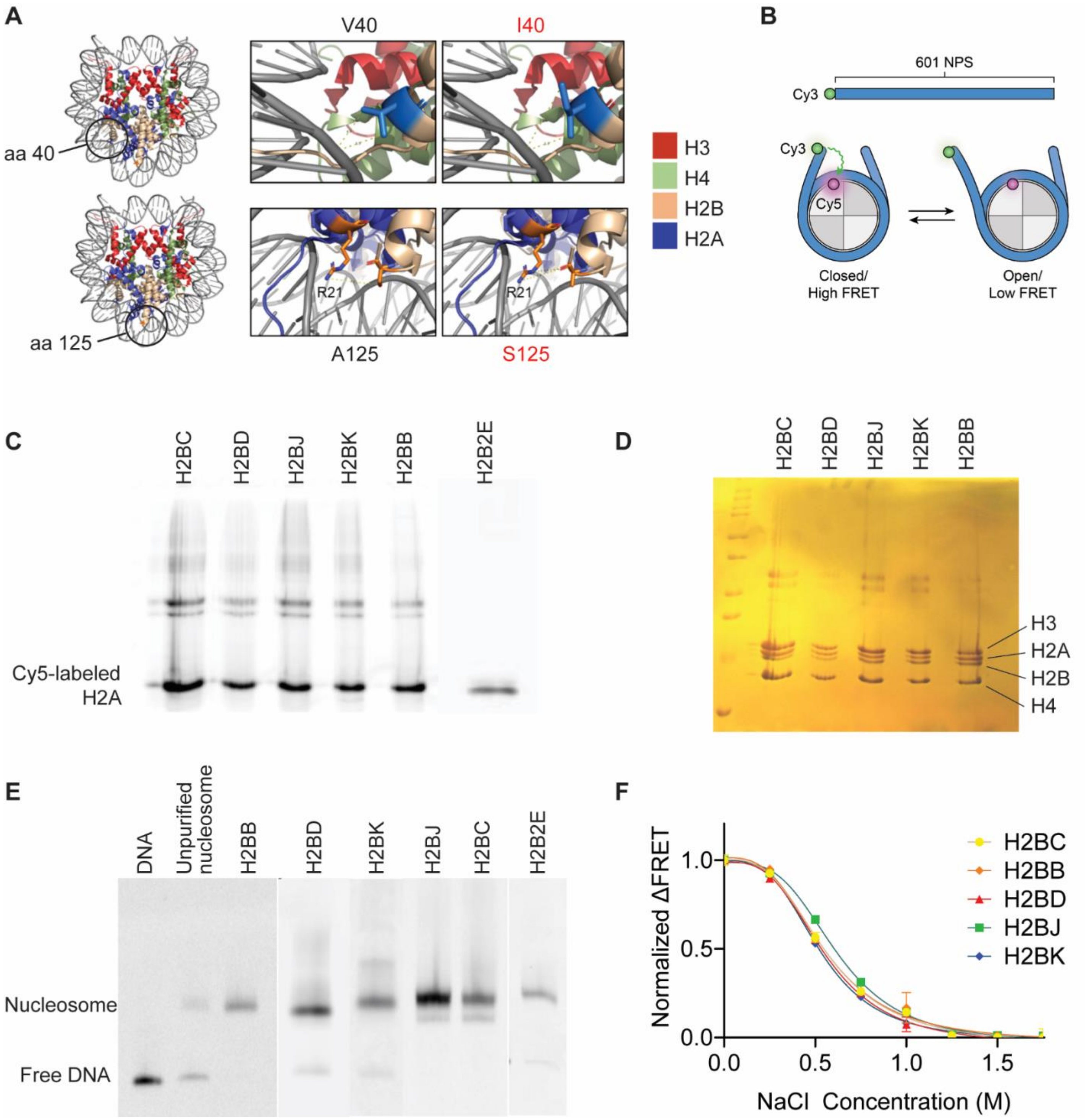


**Suppl Fig 6**. **H2B variants alter nucleosome structure and DNA accessibility. A**) Left panels are the nucleosome crystal structure based on [PDB ID: 2CV5]. H2A is shown in blue, H2B in tan, H3 in red, H4 in green, and DNA in light grey. Right panels are zoomed images at amino acid 40 in the histone fold domain (HFD), and amino acid 125 in the C-terminal tail. Their corresponding variants are highlighted in blue and orange inside the black circles, respectively. All pictures were generated using PyMOL. **B**) Schematic representation of the constructs used for *in vitro* nucleosome reconstitution. Cy5 was attached to both copies of H2AK119C. **C**) Fluorescent PAGE analysis of Cy5-labeled nucleosomes containing H2B variants. **D**) SDS PAGE stained with Coomassie blue to confirm and visualize all histones within the reconstituted nucleosomes. **E**) Native PAGE analysis of reconstituted nucleosomes. Note H2B2E was included in FRET analysis with Gal4, and not the initial NaCl-induced FRET **F**) Change in normalized FRET for canonical and variant containing nucleosomes with increasing salt concentration.

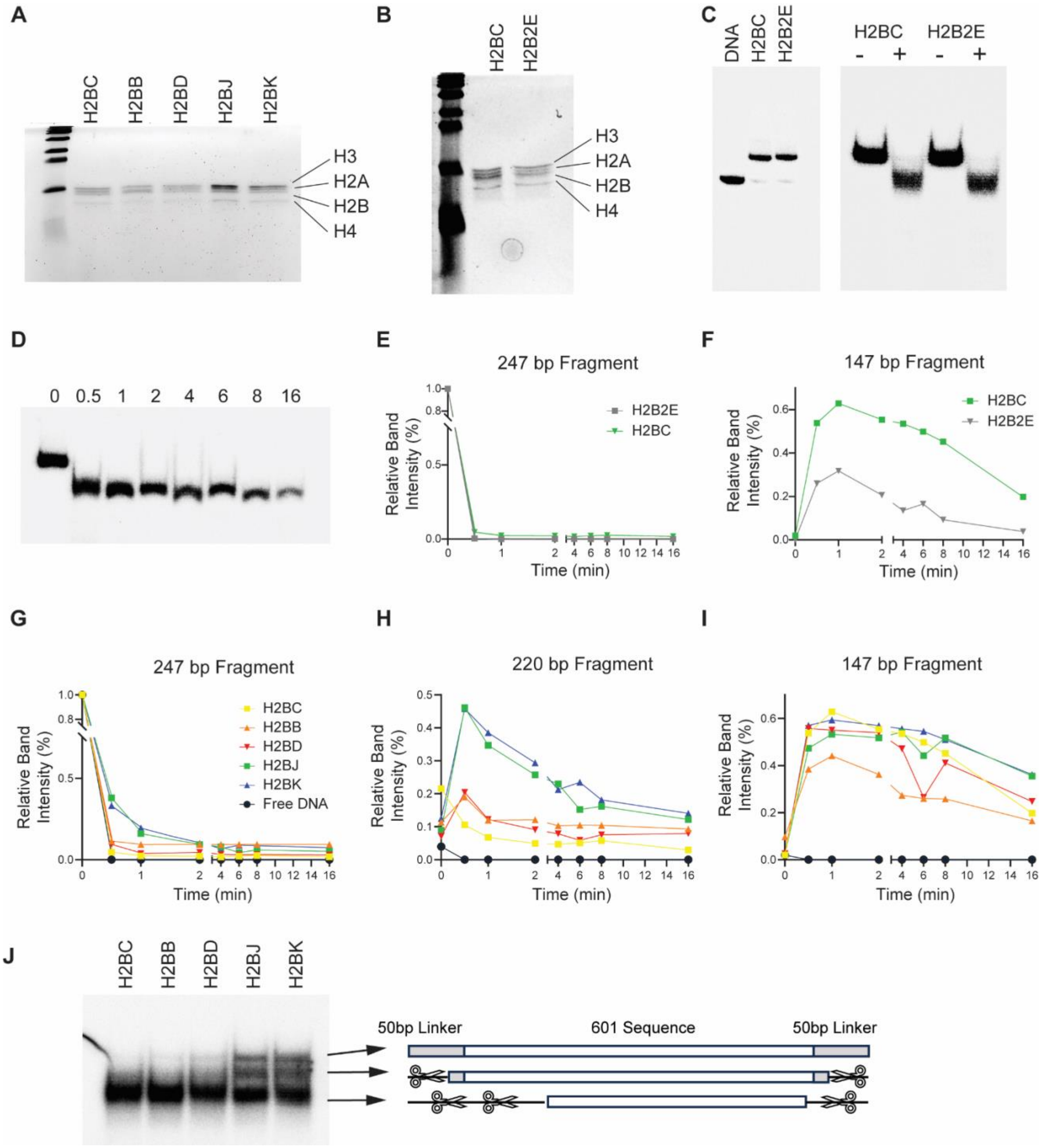


**Suppl Fig 7. Histone H2B variants alter access to nucleosome linker DNA. A-B**) SDS PAGE stained with Coomassie blue to confirm and visualize all histones within the reconstituted nucleosomes. **C**) Representative phosphor images of native PAGE analysis of naked DNA and nucleosomes used for MNase experiments. **D**) Same as **C**, except of H2B2E nucleosomes digested with MNase for the indicated times. **E-F**) Band intensities for the 247 bp (**E**) and 147 bp fragment (**F**) for the image shown in panel **D**. The intensity of each band was normalized to the 247 bp fragment at time zero. **G-I**) Band intensities for the 247 bp (**G**), 220 bp (**H**), and 147 bp (**I**) MNase digestion products for nucleosomes containing H2B variants. **J**) Native PAGE for H2B variant nucloeomes treated with for 30s with MNase. It is same image as in Fig 3C, but these digestion products (bands) were isolated from the gel, cloned, and sequenced to identify the size of each band, and the location of each cut. Digestion patterns are illustrated in the right panel. Gray boxes represent the 50 bp linker on both sides of the nucleosome. The scissor icons and dashed lines indicate where MNase cut the linker DNA. Digestion is more extensive on one side of the nucleosome than the other, indicating asymmetric accessibility.

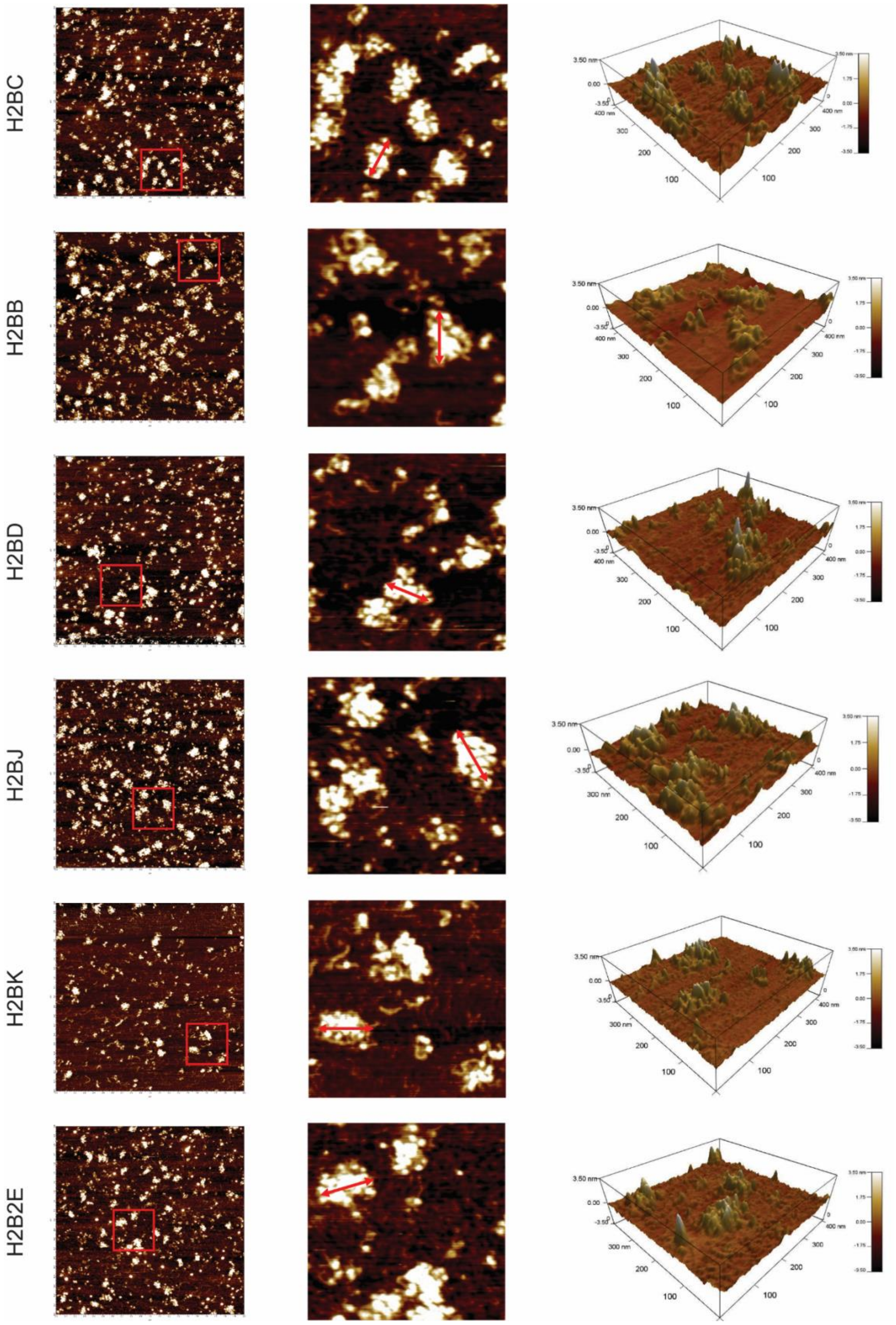


**Suppl Fig 8**. **H2B variant nucleosomes form organized nucleosome arrays.** Representative atomic force microscopy images of reconstituted chromatin arrays in 150 mM NaCl (n=3). Each red box is 400 nm$^2$ and is enlarged in the middle column. The double red arrow indicates how array diameters are measured. The right column shows the 3-dimensional topology of chromatin arrays shown in the middle column.

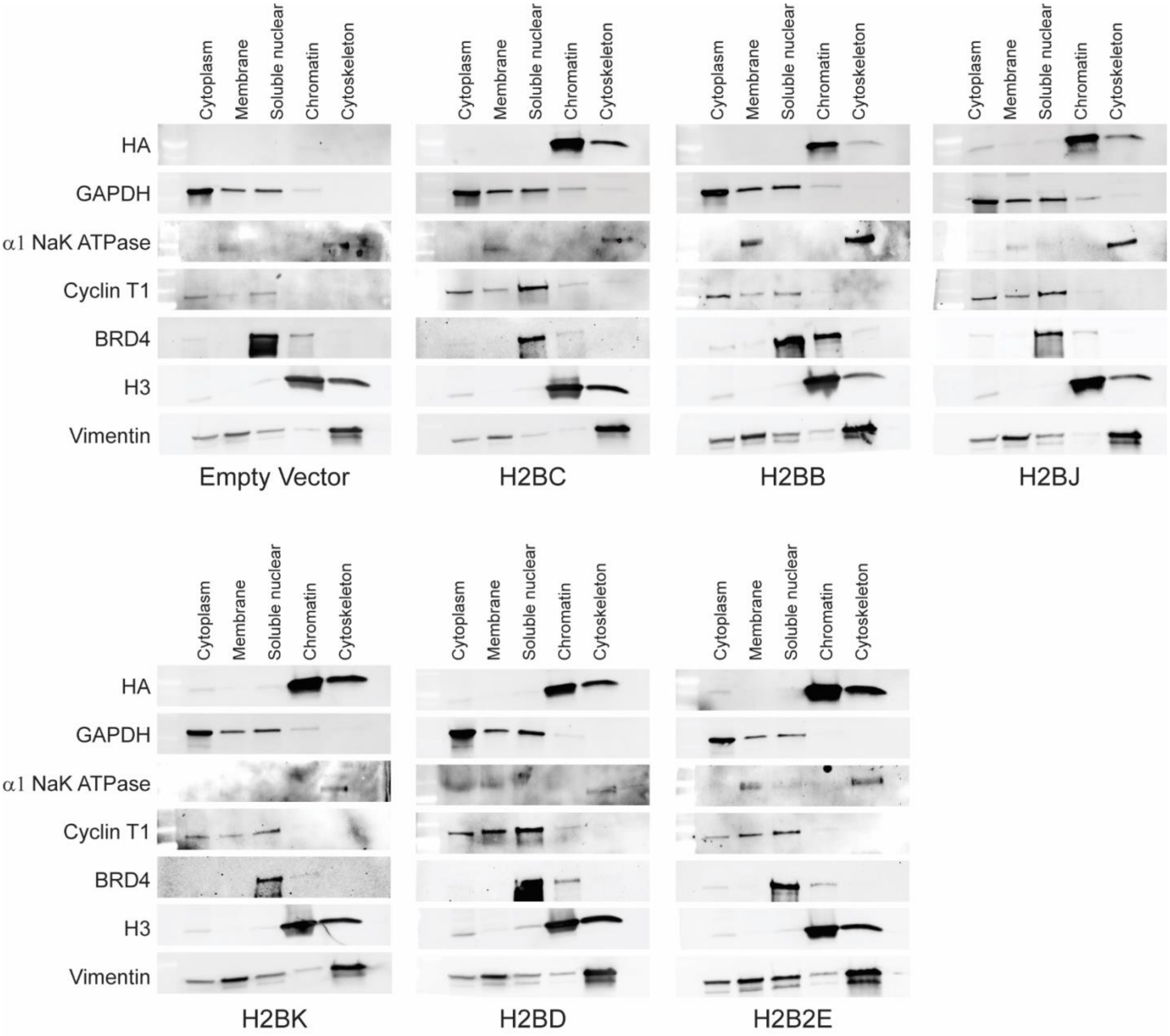


**Suppl Fig 9. Histone H2B variants are expressed and incorporated into chromatin**. Representative western blots showing *in vitro* expression of HA-tagged H2B variants in BEAS-2B cells. H3 expression served as an internal, positive control. Known localization positive controls were: GAPDH = cytoplasm; α1 NaK ATPase = membrane; Cyclin T1 and BRD4 = soluble nuclear; H3 = chromatin; Vimentin = cytoskeleton.

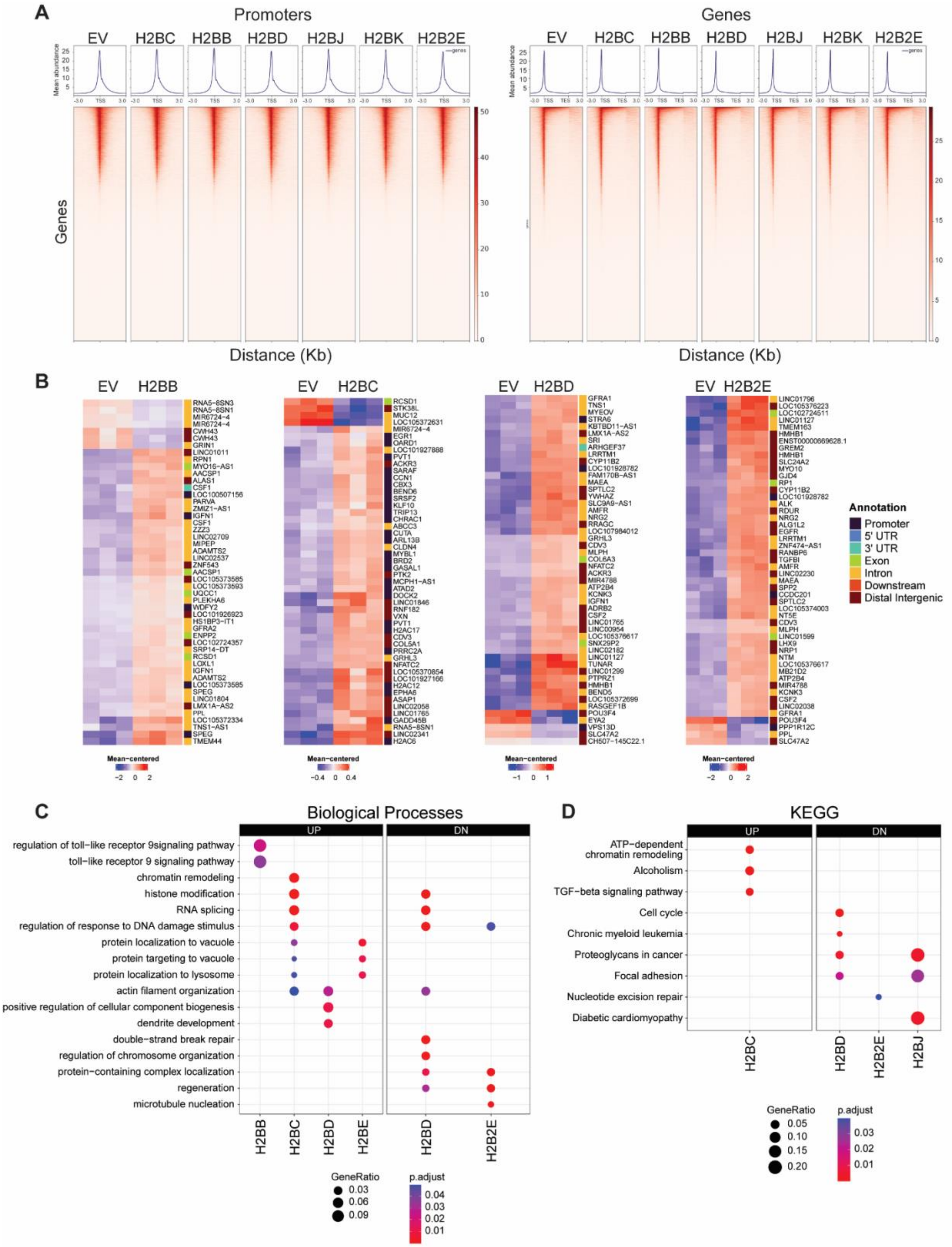


**Suppl Fig 10. Histone H2B variants alter chromatin accessibility. A**) ATAC peak proximal to gene promoters and gene bodies for BEAS-2B cells expressing each H2B variant. Only those regions within ± 3 kb of the transcription start site (TSS) are shown. EV = empty vector control. **B**) Heatmap of top 50 differentially accessible regions (DARs) in BEAS-2B cells expressing each H2B variant. DARs were defined by FDR <0.1 and Shrunken Log2(FC) > 1 for opened DARs, and Shrunken Log2FC < -1 for closed DARs. **C-D**) Dot plots of the top enriched biological processes (**C**) and KEGG (**D**) pathways in opened (UP) or closed (DN) DARs in BEAS-2B cells expressing H2B variant.

Suppl. Fig 11

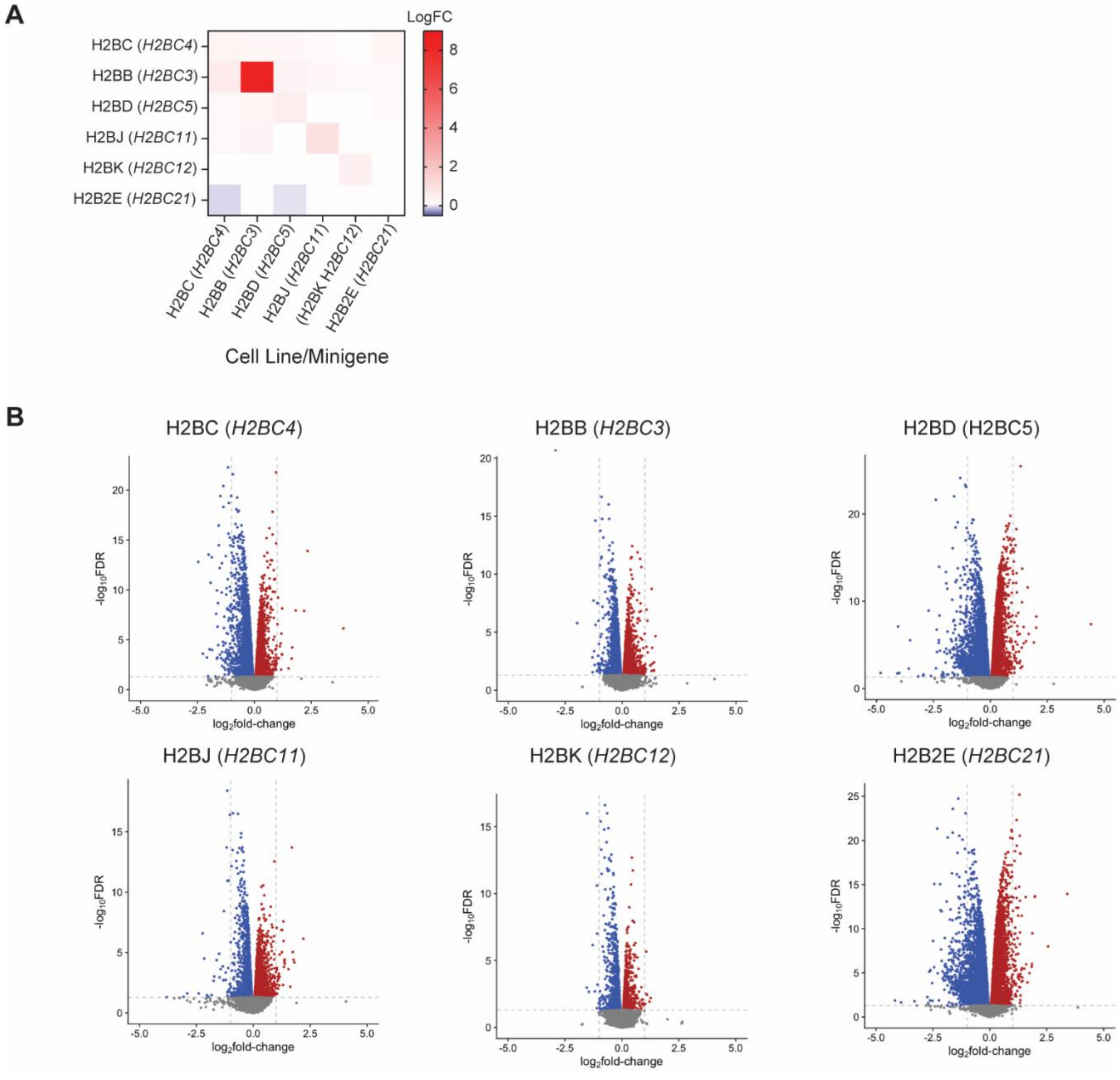
A
LogFC
8
6
4
2
0
H2BC (H2BC4)
H2BB (H2BC3)
H2BD (H2BC5)
H2BJ (H2BC11)
H2BK (H2BC12)
H2B2E (H2BC21)
H2BC (H2BC4)
H2BB (H2BC3)
H2BD (H2BC5)
H2BJ (H2BC11)
(H2BK H2BC12)
H2B2E (H2BC21)
Cell Line/Minigene
B
H2BC (H2BC4)
H2BB (H2BC3)
H2BD (H2BC5)
H2BJ (H2BC11)
H2BK (H2BC12)
H2B2E (H2BC21)
-log10FDR
log2fold-change

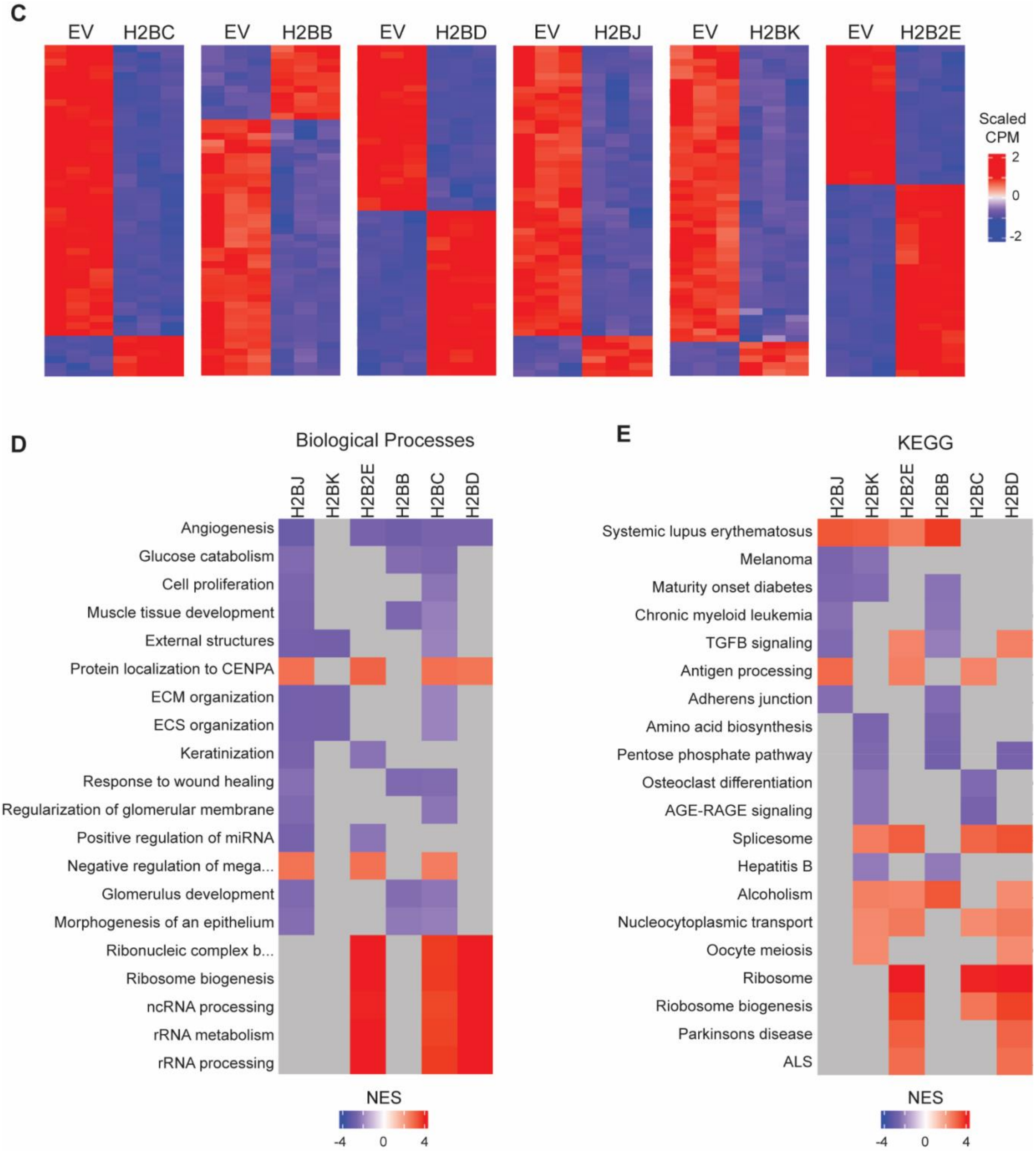


**Suppl Fig 11. Histone H2B variants alter gene expression programs. A**) H2B variant gene expression in BEAS-2B cells transfected with the indicated H2B variant minigene. Only the H2BB minigene (*H2BC3*) seemed to be overexpressed, most likely because of the low basal *H2BC3* expression from the chromosomal copy. **B**) Volcano plots of differentially expressed genes (DEGs) after incorporating of H2B variants into chromatin. BEAS-2B cells containing an empty vector were the negative control. Red = up-regulated genes; Blue = down-regulated genes; Gray = not significantly different. **C**) Top 50 DEGs for BEAS-2B cells expressing each H2B variant. **D-E**) Heatmaps of the top 20 enriched (**D**) biological processes and (**E**) KEGG pathways for H2B variant DEGs. NES = normalized enrichment score. Gray = no hits for the associated pathway.

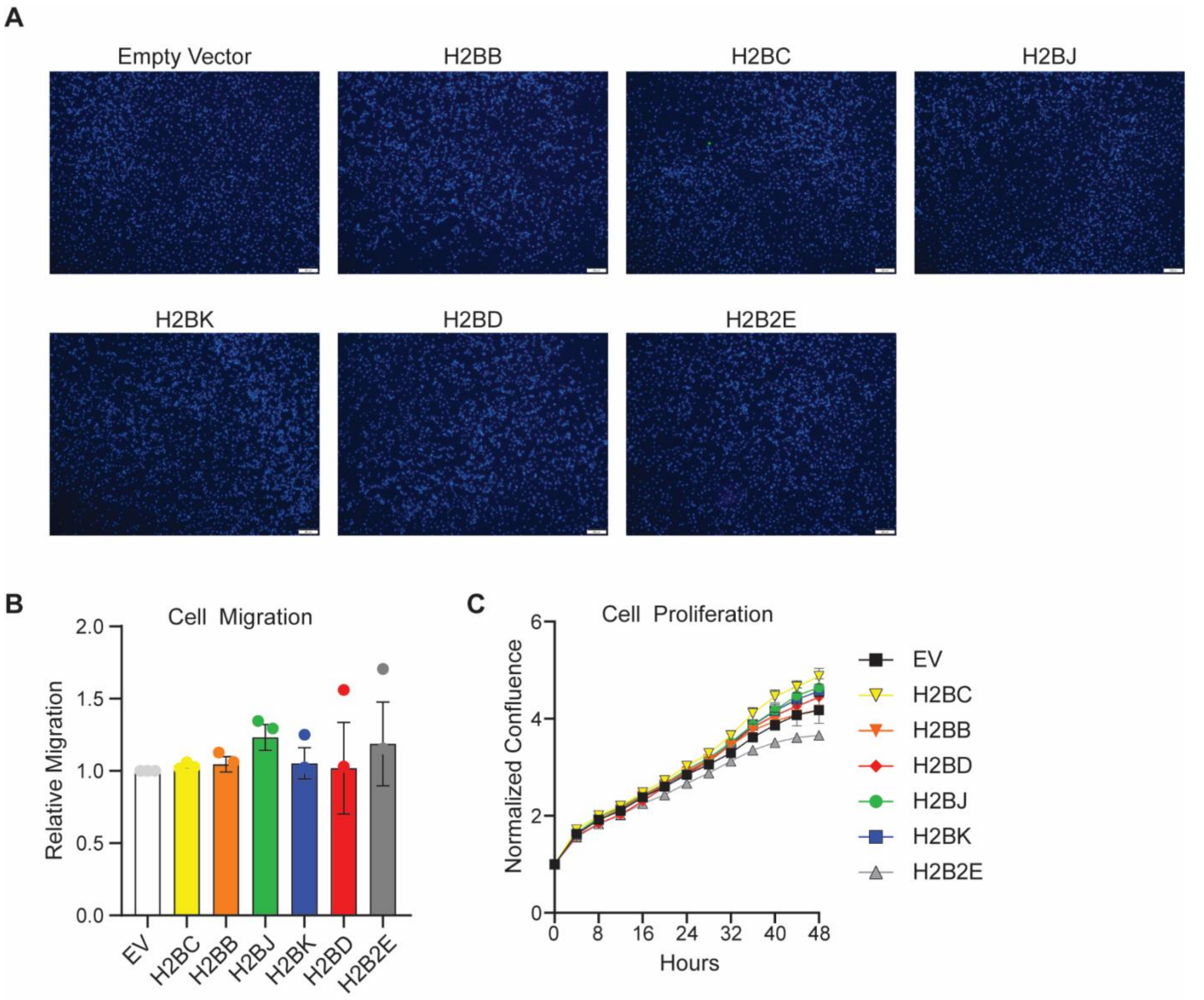


**Suppl Fig 12**. **H2B variants (by themselves) do not significantly alter cell migration or proliferation. A**) Representative images of migrating cells after 24 h in a transwell assay. Cells were fixed and stained with DAPI. **B**) Cell counts were obtained from images (like those in **A**) with an automated Olympus cellSens counting module, and normalized to BEAS-2B cells containing an empty vector (EV). **C**) Transfected BEAS-2B cell proliferation, as measured with an Incucyte assay. Cells were seeded at equal numbers and monitored through continuous image acquisition.

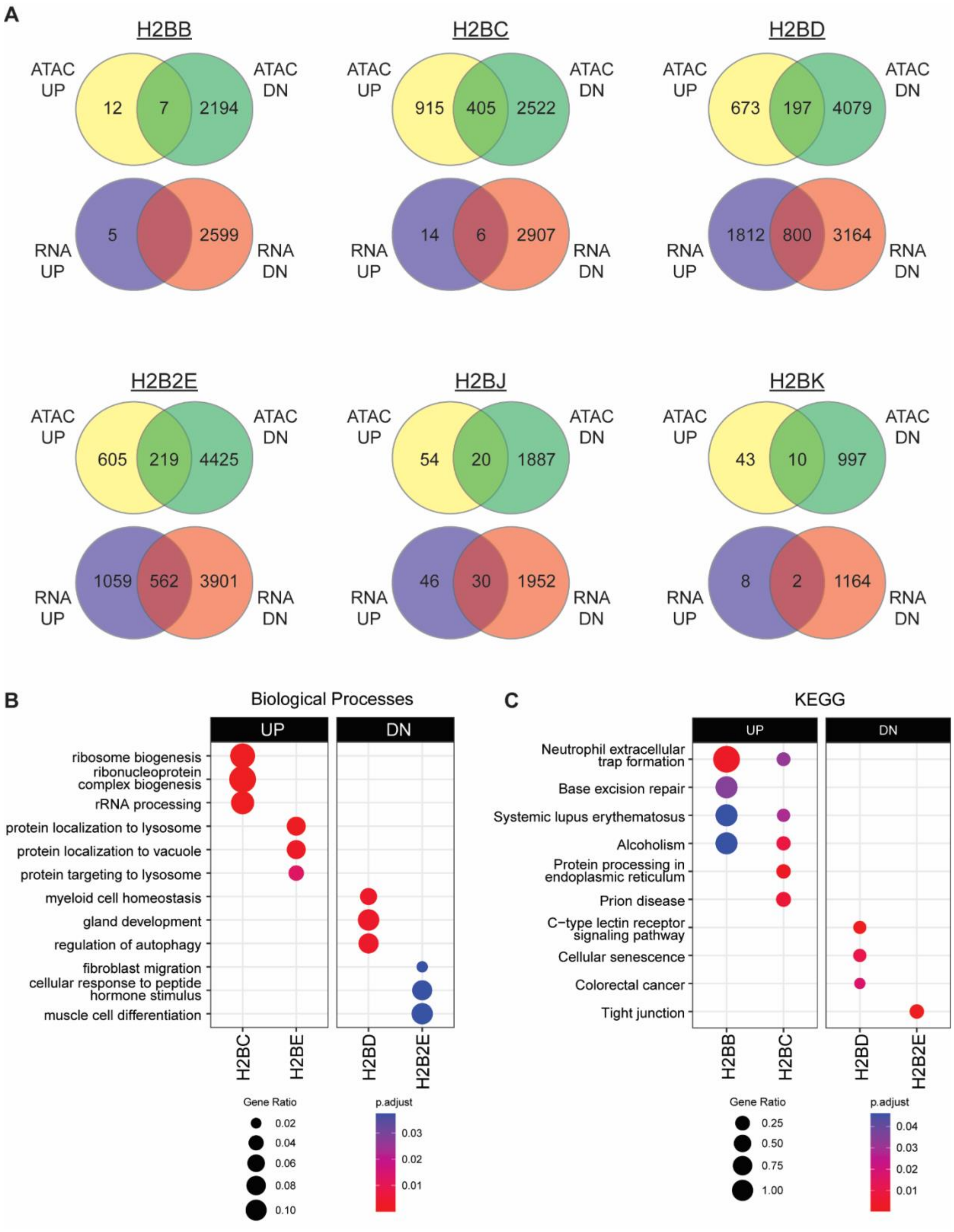


**Suppl Fig 13. H2B variant DEGs and their association with DARs. A**) Venn diagrams showing the number and overlap of opened (ATAC UP) and closed (ATAC DN) DARs relative to up- (RNA UP) and down (RNA DN)-regulated genes. **B-C**) Dotplot of the top enriched **B**) biological processes and **C**) KEGG pathways where opened (UP) and closed (DN) DARs correspond to increased and decreased RNA expression of at least five overlapping genes, respectively. Genes with differentially accessible promoters were compared with differentially expressed genes at an adjusted p-value cutoff of 0.05.
